# Detection of pathogens and antimicrobial resistant genes from urine within 5 hours using Nanopore sequencing

**DOI:** 10.1101/2024.03.04.582689

**Authors:** Anurag Basavaraj Bellankimath, Crystal Chapagain, Sverre Branders, Jawad Ali, Robert C Wilson, Truls E. Bjerklund Johansen, Rafi Ahmad

## Abstract

**Purpose:** Urinary Tract Infection (UTI) is a prevalent global health concern accounting for 1-3% of primary healthcare visits. The current methods for UTI diagnosis have a high turnaround time of 3-5 days for pathogen identification and susceptibility testing. This work is a proof-of-concept study aimed at determining the detection limit by establishing a culture and amplification-free DNA extraction methodology from spiked urine samples followed by real-time Nanopore sequencing and data analysis.

**Methods:** This study first establishes an optical density culture-based method for spiking healthy urine samples with the six most prevalent uropathogens. Pathogens were spiked at two clinically significant concentrations of 10^3^ and 10^5^ CFU/ml. Three commercial DNA extraction kits were investigated based on the quantity of isolated DNA, average processing time, elution volume and the average cost incurred per extraction. The outperforming kit was used for direct DNA extraction and subsequent sequencing on MinION and Flongle flowcells.

**Results:** The Blood and Tissue kit outperformed the other kits. All pathogens were identified at a concentration of 10^5^ CFU/ml within ten minutes, and the corresponding AMR genes were detected within three hours of the sequencing start. The overall turnaround time including the DNA extraction and sequencing steps was five hours. Moreover, we also demonstrate that the identification of some pathogens and antibiotic-resistance genes was possible at a spike concentration of 10^3^ CFU/mL.

**Conclusion:** This study shows great promise toward reducing the time required for making an informed antibiotic administration from approximately 48 hours to five hours thereby reducing the number of empirical doses and saving lives.

## 1 Introduction

Urinary Tract Infections (UTIs) are the second most prevalent infections worldwide, accounting for 1-3% of primary healthcare visits and contributing to around 13.7% of community-based antibiotic prescriptions (1,2). In 2019, there were 405 million cases of UTI globally, with 236,790 associated deaths, which is a 60% rise in cases and a 140% rise in deaths since 1990 (3,4). Left untreated, UTI can progress to pyelonephritis, a more severe kidney infection, with pregnant women facing heightened risks of preterm delivery and low birth weight babies (5). Pyelonephritis and uroepithelial invasion of pathogens can progress to urosepsis, with multiorgan failure and death (6). Urosepsis accounts for a quarter of all sepsis cases (7), and the urinary tract is the most common source of *Escherichia coli* (*E*. *coli*) bacteremia (8). Around 40% of hospital-acquired urinary tract infections may present as severe conditions such as pyelonephritis and urosepsis (9), posing a significant risk to immunocompromised, catheterized, and elderly patients if not promptly treated. In case of severe infections, it is of paramount importance to administer effective antibiotics as soon as possible. Due to a recent global increase in antibiotic resistance among urinary tract pathogens, fewer antibiotics are available as reliable empirical treatments. Hence, an increasing importance is placed on developing rapid diagnostic tests for initiating effective and lifesaving treatment as soon as possible.

UTIs are mainly caused by gastrointestinal bacteria, while fungal infections are rare. The most common pathogens include *E. coli*, *Klebsiella pneumoniae* (*K. pneumoniae*), *Proteus mirabilis* (*P. mirabilis*), *Pseudomonas aeruginosa* (*P. aeruginosa*), *Enterococcus* spp., *Staphylococcus* spp. and *Candida* spp (6,10). By consensus, a bacterial concentration of ≥10^5^ CFU/mL is the microbiological criterion for a symptomatic UTI (7). Still, depending on the clinical type and complexity, the significant bacteriuria concentrations range from 10^3^ to 10^5^ CFU/mL (11). The current diagnostic methods usually need 24 hrs to identify the pathogen, with an additional 24 hours for determining the antibiotic susceptibility of the pathogen (12). Currently, techniques such as mass spectroscopy-based matrix-assisted laser desorption ionization-time of flight (MALDI-TOF) are routinely used for pathogen identification, along with nucleic acid-based amplification tests that can perform both identification and genotypic resistance detection (13,14). While MALDI-TOF can offer rapid microbial identification, it requires prior cultivation of the pathogens. It is also difficult to differentiate closely related bacterial species, thereby reducing identification specificity (15). Meanwhile, conventional nucleic acid-based tests such as PCR are restricted to specific pathogens and lack unbiased comprehensive identification capabilities (14).

Metagenomic next-generation sequencing (mNGS) has the potential for rapid unbiased identification of pathogens at the strain level and their corresponding antimicrobial resistance genes, which overcome the limitations of the current diagnostics methods (16). mNGS is also superior for its ability to identify microbes that are difficult to culture and detect AMR genes directly without needing a targeted assay. While many commercially available sequencing technologies exist, the MinION sequencer (Oxford Nanopore Technologies, UK) exhibits excellent potential for rapid point-of-care testing. This is a pocket-sized USB-connectable, portable device capable of real-time long-read sequencing and data analysis (16,17). Previous studies have shown the feasibility of using MinION to detect bacterial pathogens and antimicrobial resistance genes from urine samples (16,18–20). Schmidt et al. detected pathogens and their corresponding antibiotic-resistance genes (ARGs) among clinical and spiked urine samples with a turnaround time of 4 - 6 hours (19). Similarly, Zhang et al. applied a metagenomic nanopore sequencing-based pipeline on 76 patient samples with a detection sensitivity of 86.7% (95% CI) and specificity of 96.8% (95% CI) compared with the conventional culture-based methods (18).

One of the critical steps for metagenomics-based diagnostic methods is DNA extraction, as variation in the extraction protocol can severely affect sequencing results. While studies have been carried out to determine the extraction efficiencies of different extraction kits on samples such as stool and blood, the urinary microbiome is relatively less explored (21,22). Some of the challenges encountered when working with urine samples are limited microbial biomass compared to other samples, the presence of different salt and urine components that can interfere with the extraction, and the presence of different assay inhibitors (23). The QIAamp BiOstic Bacteremia DNA Kit, DNeasy Blood & Tissue Kit, and PowerFood Microbial Kit have been the most used kits for the isolation of DNA from urine samples. Particularly, the BiOstic Bacteremia DNA Kit and the Blood & Tissue Kit were found to preserve the bacterial diversity of the sample while yielding a large amount of good-quality DNA for downstream sequencing (21,24,25).

The current work is a proof-of-concept study aimed at determining the detection limit by establishing a culture and amplification-independent methodology of extracting DNA from spiked urine samples followed by Nanopore sequencing on MinION and Flongle flowcells. To achieve this, we first established an optical density (OD) based culture method for preparing spiked urine samples at clinically relevant concentrations of 10^5^ and 10^3^ CFU/mL using the most common uropathogens based on their prevalence in UTI cases. This was followed by benchmarking of three commercial DNA extraction kits: QIAamp BiOstic Bacteremia DNA Kit, DNeasy Blood & Tissue Kit, and PowerFood Microbial Kit. The blood and tissue kit outperformed the other two based on the quantity of isolated DNA, average processing time, elution volume and the average cost incurred per extraction. The extracted DNA was then sequenced on nanopore platforms with the subsequent characterization of the pathogens and their corresponding ARGs. All the pathogens and their corresponding antibiotic-resistant genes at a spike concentration of 10^5^ CFU/mL were identified between 10 to 200 minutes post-sequencing. The method including all the extraction, sequencing and bioinformatic analysis steps has an overall Turn-Around-Time (TAT) of 5 hours. Additionally, three spiked pathogens *E. coli* NCTC13441, *K. pneumoniae* CCUG225T and *P. mirabilis* CCUG2676T were identified at 10^3^ CFU/mL indicating the method potential at CFUs lower than 10^5^. Such a culture-independent workflow that combines real-time sequencing and data analysis could be transformative for clinical management of severe UTIs including urosepsis.

## 2 Methods

### 2.1 Ethics statement

Human urine samples were obtained from anonymous healthy donors at the Department of Biotechnology, Inland Norway University of Applied Sciences (INN). Urine samples were spiked with different bacterial strains, as mentioned below. In line with our previous work (26), there was no intention to sequence human DNA. Therefore, commercial kits for bacterial DNA extraction were used. Moreover, any sequencing reads recognized as being generated from human DNA were omitted from further analysis and permanently discarded. The microbiology laboratory at INN is approved for the described experimental work. All methods were carried out in accordance with relevant guidelines and regulations of the INN University.

### 2.2 Experimental design

#### 2.2.1 Inoculum optimization of the bacterial strains relevant to UTI

The current study used six clinically relevant bacterial species commonly found in UTIs, including *Escherichia coli, K. pneumoniae, P. mirabilis, E. faecalis, P. aeruginosa & S. aureus,* (Supplementary Table I) as test strains. Multiple strains of *E. coli* were included in the study, as most UTI cases are associated with *E. coli* species. Two E. coli isolates (*E. coli NCTC 13441 & INN A2-39*) and the *K. pneumoniae* isolates were positive for the presence of extended-spectrum β-lactamases (ESBL). The strains *K. pneumoniae* CCUG225T, *E. faecalis* CCUG 9997 (R), and P. aeruginosa CCUG17619 (R) were retrieved from the culture collection at the University of Gothenburg (CCUG, Sweden). The strains *E. coli* NCTC 13441 (R) and methicillin-sensitive strain *S. aureus* NCTC8325 were retrieved from the National Collection of Type Cultures (NCTC, UK Health Security Agency, UKHSA, UK). Additionally, three clinical strains (INN 125, INN A2-39 & INN 155) were sourced from our in-house collection at INN University, Hamar.

The bacterial strains were revived by streaking a loopful of frozen glycerol stock on Brain Heart Infusion (BHI) agar plates (15 g/L Agar, 37 g/L BHI Broth, VWR Life Science, USA) and incubated at 37±2 °C overnight. Individual colonies from the overnight culture were resuspended in 1X PBS buffer saline (E504-100ML, VWR Life Science, USA), and the optical density of each suspension was measured at 600 nm using a UV 3100PC Spectrophotometer (VWR Life Science, USA). To determine the colony formation units (CFU/mL), the prepared suspension of known OD was serially diluted up to 10^-6^ dilutions, and 50 µL of 10^-4^, 10^-5^, and 10^-6^ dilutions were spread on BHI agar plates and incubated at 37±2 °C for 24 hours. Four to seven suspension replicates were prepared for all the bacterial strains used in the study for the determination of the correlation between the optical density (A600) and CFU/mL (Supplementary Table II). The optical density was used as a starting reference point and precise viable bacteria numbers were determined using the plate count method (Supplementary Figure I).

#### 2.2.2 Evaluation of DNA extraction kits

Based on the literature review, three kits, the QIAamp BiOstic Bacteremia DNA Kit (Qiagen, Germany), DNeasy Blood & Tissue Kit (Qiagen, Germany), and PowerFood Microbial Kit (Qiagen, Germany) were selected and compared in terms of DNA yield, protocol length, and cost per sample. The kits were tested on two gram-negative bacteria (*E. coli* and *K. pneumoniae*) and one gram-positive bacterium (*S. aureus*). All the pre-treatment steps, if present in the respective kit protocol, were included in the testing. Finally, the best-performing kit was selected based on the quantity of isolated DNA, average processing time, elution volume and the average cost incurred per extraction for the direct DNA extraction steps.

### 2.3 Direct DNA extraction from spiked urine samples and Sequencing

#### 2.3.1 For MinION sequencing

As presented in the inoculation optimization section, a bacterial suspension of the strain to be spiked was prepared in 1X PBS by transferring bacterial colonies from a fresh overnight-grown culture. The OD of the prepared suspension was then adjusted to correlate with either 10^9^ or 10^8^ CFU/mL, depending on the target bacterial species (Supplementary Table II). The CFU of the prepared suspension was verified by the standard spread plate method. Only the spiked samples verified to have the target 10^9^ or 10^8^ CFU/mL were used further in downstream steps. For preparation of spiked urine samples, a 30 ml clean catch midstream urine sample from a healthy donor was collected in a sterile bottle as described by Cheesbrough et al. (27). The prepared bacterial suspension with 10^9^/10^8^ CFU/mL was used to spike 30 mL of collected urine sample to the final desired concentrations of 10^3^, 10^4^, and 10^5^ CFU/mL spiked urine. The spiked samples were then pre-processed by the addition of 10% (v/v) of 1 M Tris-EDTA (10 mM Tris, 1 mM EDTA, pH 8.0) and centrifuged at 3260 x *g* for 10 minutes at 4 °C as described by Munch M M et al., (28). The resulting pellet was used for further DNA extraction by the best-performing extraction kit determined from the **section 2.2.2**. After extraction, the concentration and purity of the DNA samples were determined using NanoDrop (Thermo Scientific, USA) and the Qubit dsDNA HS Assay kit (Thermo Fisher Scientific) and were further used for library preparation and sequencing (Figure I).

**Figure I:**
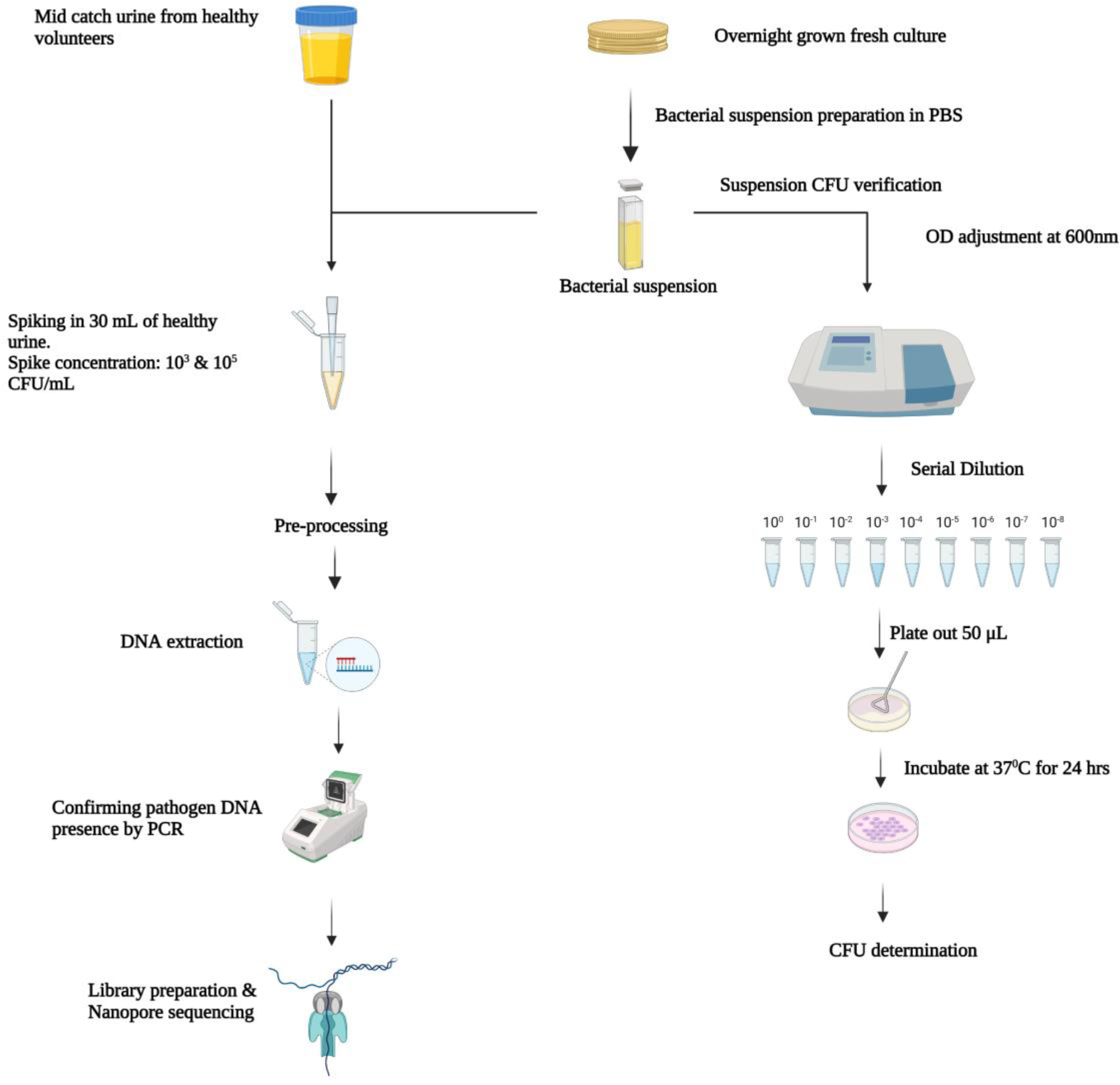
A graphical overview of the steps involved in the inoculum preparation, spiking, DNA extraction and sequencing steps involved in the study.

#### 2.3.2 For Flongle sequencing

In addition to sequencing samples on MinION flow cells, bacterial spiked samples were also sequenced on the Flongle flow cells. The rationale behind this was to check the possibility of replicating results obtained from MinION flow cells on the Flongle flow cell, a cost-effective, one-time-use version mounted on the MinION sequencing device using a Flongle adapter. Three pathogens, INN 125, *E. faecalis* CCUG 9997, and *P. aeruginosa* CCUG17619, were spiked in 5 mL of healthy urine samples to achieve 10^3^, 10^4^, and 10^5^ CFU/mL concentrations. DNA was directly extracted from the spiked samples, as described in the section **2.3**.

### 2.4 PCR-based identification of spiked pathogenic & host DNA

Before sequencing, a PCR-based verification step was performed on the extracted samples to determine the presence of pathogenic DNA. The primers used for the verification are mentioned in supplementary table III. All amplification reactions were carried out in a final volume of 20 µL of reaction mixture containing 4 µL of 5 X HOT FIREPol® MultiPlex Mix Ready to Load with 10 mM MgCl_2_ (Solis BioDyne, Estonia), 0.5 µL of 10 µM forward and reverse primers and 1-8 ng template DNA. For the negative controls, the template DNA was replaced with nucleic acid- and nuclease-free water. The PCR reactions were carried out in a Verity 96-Well Thermal Cycler (Applied Biosystems, USA), with the following thermal profile: initial denaturation for 12 min at 95 °C followed by 30 cycles of 25 s at 95 °C, 45 s at 60 °C and 45 s at 72 °C, with final extension at 72 °C for 7 min. Following PCR, the obtained products were visualized on 1% agarose gel electrophoresis (AGE) premixed with SYBR safe (Invitrogen, USA) gel DNA stain. Ten µL of PCR product was loaded onto the gel, and electrophoresis was run for 45 minutes at 80 V, whereafter the DNA fragments were visualised using G: box (SYNGENE, USA) and UV-illuminator (Invitrogen, USA).

### 2.5 Library preparation & sequencing on MinION and Flongle flow cells

#### MinION flow cell

Sequencing was performed on DNA directly extracted from urine samples spiked with bacterial concentrations of 10^3^ and 10^5^ CFU/mL. Due to the very low initial concentration of the isolated DNA, it was first concentrated and purified using the Agencourt AMPure XP system (Beckman Coulter, USA) according to the manufacturer’s instructions. The library of purified DNA samples was prepared using Rapid Barcoding Sequencing kit SQK-RBK004 (Oxford Nanopore, UK) following the manufacturer’s protocol. The adaptor-ligated library was loaded onto the MinION flow cells (R9.4.1 FLO-MIN 106D, Oxford Nanopore) for sequencing.

#### Flongle flow cells

Similar to MinION-based sequencing, the extracted DNA was first concentrated using the Agencourt AMPure XP system (Beckman Coulter, USA) according to the manufacturer’s instructions. The library was prepared using the Ligation Sequencing Kit (SQK-LSK109) and the Native Barcoding Expansion Kit (EXP-NBD104) following the manufacturer’s protocol. The prepared library was then loaded onto the Flongle flow cells (R9.4.1 FLO-FLG001, Oxford Nanopore) for sequencing.

### 2.6 Bioinformatic analysis of the sequencing data

The data from the MinION and Flongle flow cells were collected using the MinKNOW software (version 5.3.1) with the default settings, except for the sequencing run time, which was set to 18 hours in the case of MinION and at least 24 hours for Flongle. Base calling was performed in real-time using the Guppy tool (version 6.3.9) incorporated within the MinKNOW software. The Fastq files obtained after basecalling were first converted to FASTA files using BBMap’s (version 38.84) reformat.sh tool, then analysed for taxonomy classification, AMR gene detection, and reference genome coverage.

The identification of pathogens was achieved by aligning the FASTA nanopore reads using blastn (29) (version 2.14.0), against a database containing all NCBI RefSeq genomes of the spiked species. Reads shorter than 500 bp were ignored in the analysis. Hits were filtered with stringent quality criteria to avoid false positives. These criteria were sequence identity of at least 90 % and coverage of at least 70 % of the read in the alignment. Next, the reads that did not meet these criteria were BLAST searched against the NCBI prokaryotic reference (RefProk) database (30) and filtered again with the same stringent criteria. Reads that did not meet the criteria after BLAST search against the species-specific nor the RefProk database were considered not to be of prokaryotic origin.

The raw sequences in FASTA format were screened for resistance genes using the ABRicate tool (version 1.0.1) (31) with the CARD (32), ResFinder (33), MEGARES (34) and NCBI AMRFinderPlus (35) and Resfinder (33) databases with default parameters for AMR gene detection. For genome coverage analysis, the reads were aligned to their reference genome using blastn, and the percentage of total alignment was calculated based on the overall number of nucleotides covered in the alignment.

## 3 Results

### 3.1 Establishment of the optimal experimental conditions to replicate clinical UTI-relevant urine samples

Optical Density (OD) provided a preliminary estimate of the bacterial concentration for spiking the urine samples. The OD of the PBS suspension of fresh overnight bacterial culture was measured using a spectrophotometer and then cultured on BHI agar to calculate colony-forming units. The protocol optimization experiments were initially conducted using only the wild-type *E. coli* CCUG 17620 strain. A series of suspensions between OD 1.2 and 1.8 at 600 nm were prepared, corresponding to cell concentrations ranging from 2.1*10^8^ to 2.1*10^9^ CFU/mL (Supplementary Table II). Although OD provides a preliminary assessment of cell concentration, it does not necessarily correlate precisely to viable bacteria. Hence, the prepared bacterial suspensions were always plated out to determine the exact CFU/mL.

### 3.2 Comparison of DNA isolation kits

The performance of three commercial kits, BiOstic, PowerFood, and Blood and Tissue, was compared in terms of DNA yield, protocol length, and cost per sample to select the best kit for direct DNA extraction from spiked urine samples. The comparison was performed using two gram-negative bacteria (*E. coli* and *K. pneumoniae*) and a gram-positive species, *S. aureus*. The blood and Tissue kit outperformed the other two in terms of the quantity of isolated DNA, average processing time, elution volume and the average cost incurred per extraction (Table I). The PCR amplification gel images for all three strains are presented in the supplementary Figure II.

**Table I:**
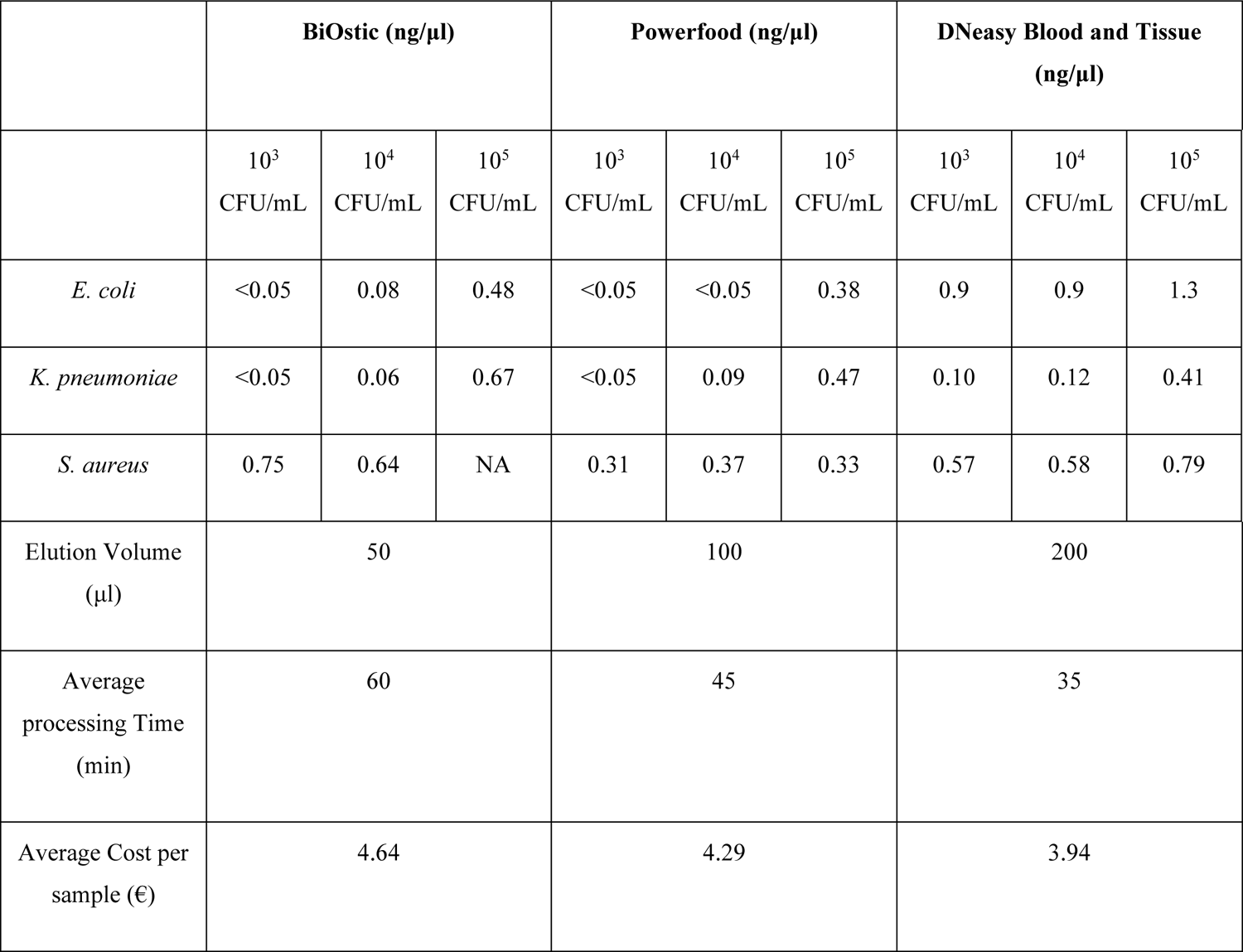
Comparison of DNA concentration isolated using three different kits. The DNA yield of BiOstic, PowerFood, and DNeasy Blood and Tissue kits was compared using two gram-negative and one gram-positive bacteria. In the case of *S. aureus*, the DNA was isolated using two different strains (*S. aureus CCUG 17621* for BiOstic and *S. aureus NCTC 8325* for PowerFood and Blood and Tissue). NA indicates no DNA was isolated.

### 3.3 Comparison of DNA yields and PCR confirmation of spiked pathogenic DNA

DNA was isolated from the urine samples spiked with different concentrations of five *E. coli* strains, three *S. aureus* strains, and one strain each of *K. pneumoniae*, *P. mirabilis*, *E. faecalis*, and *P. aeruginosa* using DNeasy Blood and Tissue kit. The average concentration of DNA isolated from the urine samples spiked with 10^3^ and 10^5^ CFU/mL bacteria was less than 1 ng/μL (0.05 ng/μL to 0.9 ng/μL) (Figure II). Remarkably, the concentration of DNA isolated from samples having the same CFU/mL of different bacteria varied widely, with the highest yield from *E. coli* and the lowest from *E. faecalis.* Furthermore, the amount of DNA extracted increased gradually from cultures of the same bacterial species with increasing concentrations of 10^3^ CFU/mL to 10^5^ CFU/mL. *E. faecalis* was an exception, which exhibited lower DNA concentration at 10^5^ CFU/mL than at 10^3^ CFU/mL.

**Figure II:**
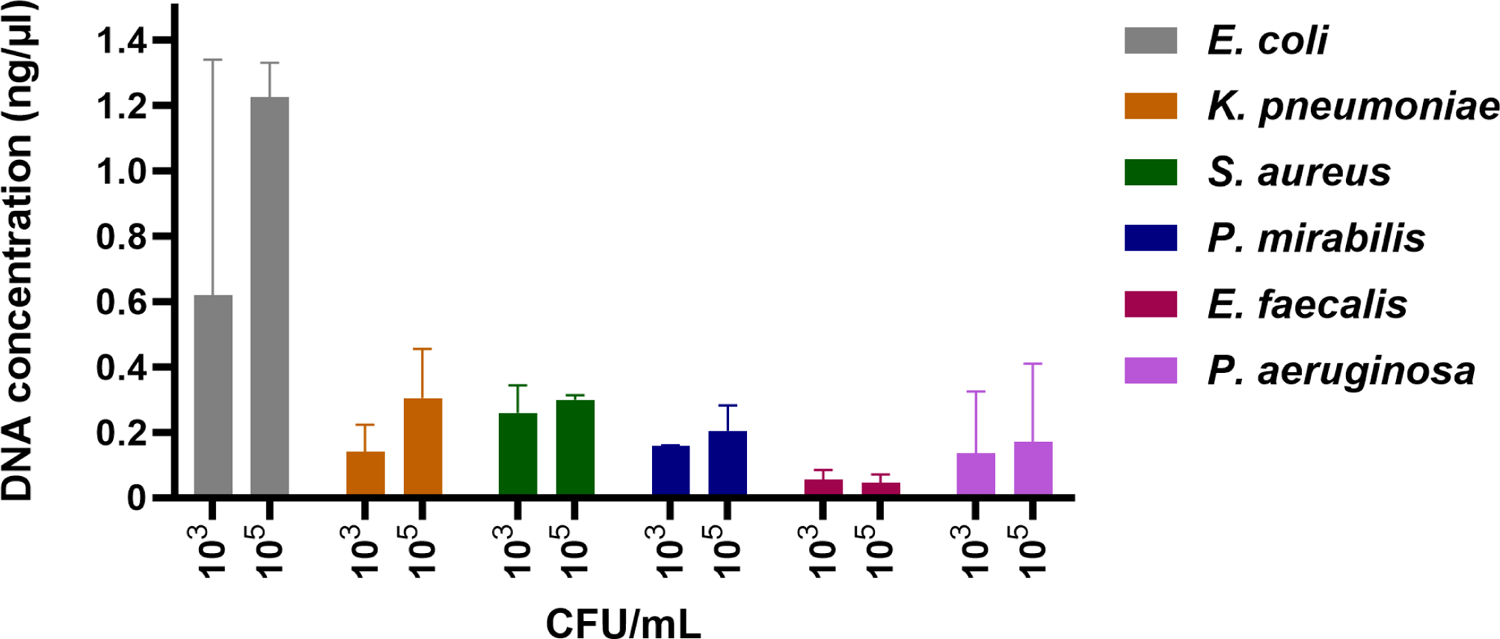
Comparison of DNA concentration among different species. The DNA was extracted from 30 ml urine samples spiked with 10^3^ and 10^5^ CFU/mL of bacteria using a Blood and Tissue kit. The DNA concentration was measured using Qubit.

To confirm the presence of spiked bacterial DNA in the extracted samples, species-specific PCR amplification was performed. The samples spiked with 10^3^ and 10^5^ CFU/mL were PCR-positive for all the species (Supplementary Figure II & Supplementary Figure III).

### 3.4 Direct Nanopore Sequencing using MinION Flowcells

The presence of pathogens and antimicrobial resistance genes (ARGs) from urine samples spiked with different clinically relevant bacterial species at two different concentrations, 10^3^ and 10^5^ CFU/mL, was investigated using the MinION platform. The raw data was base called in real-time using the integrated base caller Guppy within the MinKNOW software. The first 4000 reads generated on MinION flow cells from each spiked sample at the 10^3^ & 10^5^ CFU/mL concentrations were used as queries in nBLAST searches against the NCBI RefSeq database. The obtained results for six prevalent UTI pathogens *E. coli NCTC13441*, *K. pneumoniae* CCUG225T, *P. aeruginosa* CCUG17619, *S. aureus* NCTC8325, *E. faecalis* CCUG9997 and *P. mirabilis* CCUG2676T are summarised in the Table II & Figure III. The results showed that all pathogens were successfully identified at 10^5^ CFU/mL concentrations. In comparison, at 10^3^ CFU/mL, pathogen identification was possible from three out of nine samples spiked. The results for the three internal strains INN 125, INN A2-39 and INN 155 are presented in Supplementary Figure IV.

**Figure III:**
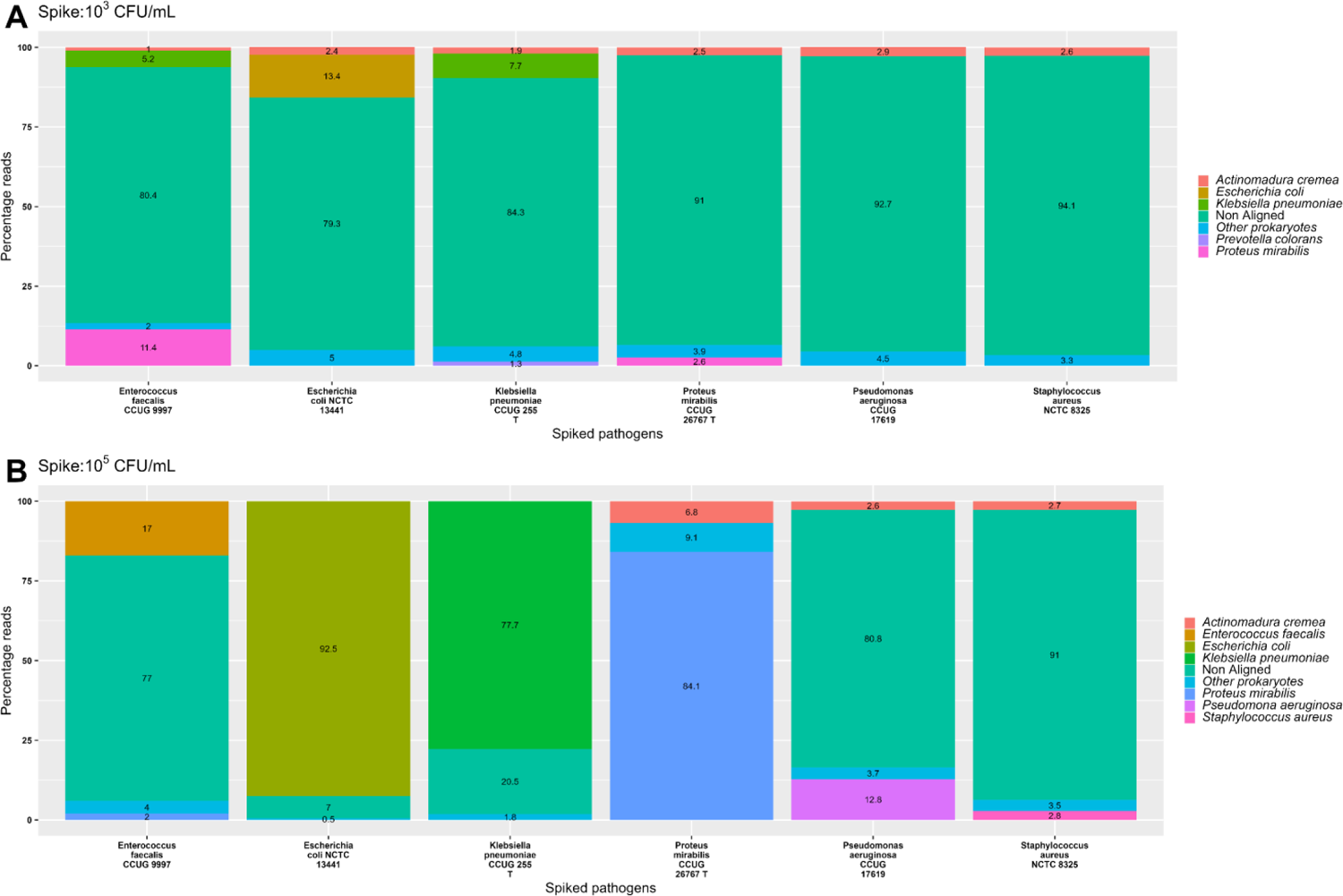
The figure denotes the distribution of reads from the sequence data generated by nanopore sequencing of extracted DNA from spiked urine samples on the MinION flow cell. The tabulated results are from the first 4000 reads generated by the MinION platform within 10 minutes (approximately) of the start of sequencing. The subfigure A & B represent the samples spiked at 103 and 105 CFU/mL, respectively. The denoted percentage reads are based on the BLAST search against the RefProk database (prokaryotic sequence data only). The data for the rest of the three in-house strains (INN A2-39, INN 125, and INN 155) is presented in Supplementary Figure III.

**Table II:** Summary of the Pathogen Identification and AMR gene detection by MinION nanopore sequencing from two different concentrations of spiked urine samples. The order of the organisms is based on their prevalence in complicated UTIs. Green ticks indicate detected, and red ticks indicate not detected for both pathogen identity and Antibiotic Resistant Genes (ARG).

| Strains | CFU/mL | DNA (ng/μl) | Pathogen Id | ARG |
| --- | --- | --- | --- | --- |
| *E. coli* NCTC13441 | $10^3$ | 0.90 | ✓ | ✓ |
| | $10^5$ | 1.30 | ✓ | ✓ |
| INN 125 | $10^3$ | 1.13 | ✗ | ✗ |
| | $10^5$ | 1.15 | ✓ | ✗ |
| INN A2-39 | $10^3$ | 0.88 | ✗ | ✗ |
| | $10^5$ | 0.97 | ✓ | ✓ |
| INN 155 | $10^3$ | 0.05 | ✗ | ✗ |
| | $10^5$ | 0.20 | ✓ | ✓ |
| *E. faecalis* CCUG9997 | $10^3$ | 0.08 | ✗ | ✗ |
| | $10^5$ | 0.24 | ✓ | ✗ |
| *K. pneumoniae* CCUG225T | $10^3$ | 0.10 | ✓ | ✗ |
| | $10^5$ | 0.41 | ✓ | ✓ |
| *P. mirabilis* CCUG2676T | $10^3$ | 0.15 | ✓ | ✗ |
|  | 10<sup>5</sup> | 0.26 | ✓ | ✓ |
| ***P. aeruginosa* CCUG17619** | 10<sup>3</sup> | 0.27 | ✗ | ✗ |
|  | 10<sup>5</sup> | 0.34 | ✓ | ✓ |
| ***S. aureus* NCTC 8325** | 10<sup>3</sup> | 0.57 | ✗ | ✗ |
|  | 10<sup>5</sup> | 0.79 | ✓ | ✓ |

The criteria for taxonomic identification of the spiked pathogen in a sample was set to be the highest number of bacterial reads in the taxonomic classification. The results also indicate the presence of 1-3% of reads labelled as *Actinomadura cremea* in a few samples at both concentrations. The remaining bacterial species (excluding the spiked pathogen) that were identified and had less than 1% assigned reads were collectively presented as other prokaryotes.

In the sample spiked with 10^5^ CFU/mL of *E. coli NCTC 13441*, 92.5% of the reads were assigned to E. coli, 7% were non-aligned, and 0.5% were other prokaryotes. Compared to the sample spiked with 10^3^ CFU/mL of the same bacteria, the percentage of non-aligned reads increased from 7% to 80%, and the percentage of bacterial reads changed from 92.5% to 13%. Similar results were also observed in samples spiked at 10^3^ & 10^5^ CFU/mL concentrations of *K. pneumoniae CCUG225T* and *P. mirabilis CCUG2676T*. However, in all cases, the samples exhibited the highest number of prokaryotic reads assigned to the respective spiked pathogen, which confirms the pathogen identification at both CFU/mL concentrations for *E. coli NCTC 13441, K. pneumoniae CCUG225T* and *P. mirabilis CCUG2676T*.

For samples spiked with *P. aeruginosa* CCUG 17619 and *S. aureus* NCTC 8325, the percentage of non-aligned reads was 92.7% & 94.1% at 10^3^ CFU/mL and decreased in samples spiked at 10^5^ CFU/mL concentration to 80.8% and 91%, respectively. While 12.8% & 2.8% of the respective reads at 10^5^ CFU/mL were assigned to *P. aeruginosa* and *S. aureus*, no spiked pathogen reads were detected at 10^3^ CFU/mL. Hence, the taxonomic identification was established only at 10^5^ CFU/mL spike concentration for these two pathogens.

However, the sample spiked with 10^3^ CFU/mL *E. faecalis* CCUG9997 was misclassified. The obtained reads comprised 79% non-aligned, 12% *P. mirabilis*, 6% *K. pneumoniae,* and 3% other prokaryotes. On the other hand, the sample spiked with 10^5^ CFU/mL *E. faecalis* CCUG9997 contained 77% non-aligned reads, 16% *E. faecalis*, 3% *P. mirabilis*, 1% *Actinomadura cremea*, and 3% other prokaryotes. Consequently, the target pathogen, *E. faecalis*, was only identified in the sample spiked with 10^5^ CFU/mL.

#### 3.4.1 Detection of antibiotic resistance genes

The ABRicate tool was used to search antibiotic resistance genes in four different databases (CARD, MEGARES, ResFinder, and NCBI AMRFinderPlus), combining the obtained results during the analysis stage. Of the nine samples spiked with 10^5^ CFU/mL of the pathogenic bacteria, reference AMR genes were detected in seven, while in samples spiked with 10^3^ CFU/mL of the pathogenic bacteria, the detection was possible only in the sample spiked with *E. coli* NCTC 13441 (Figure IV).

**Figure IV:**
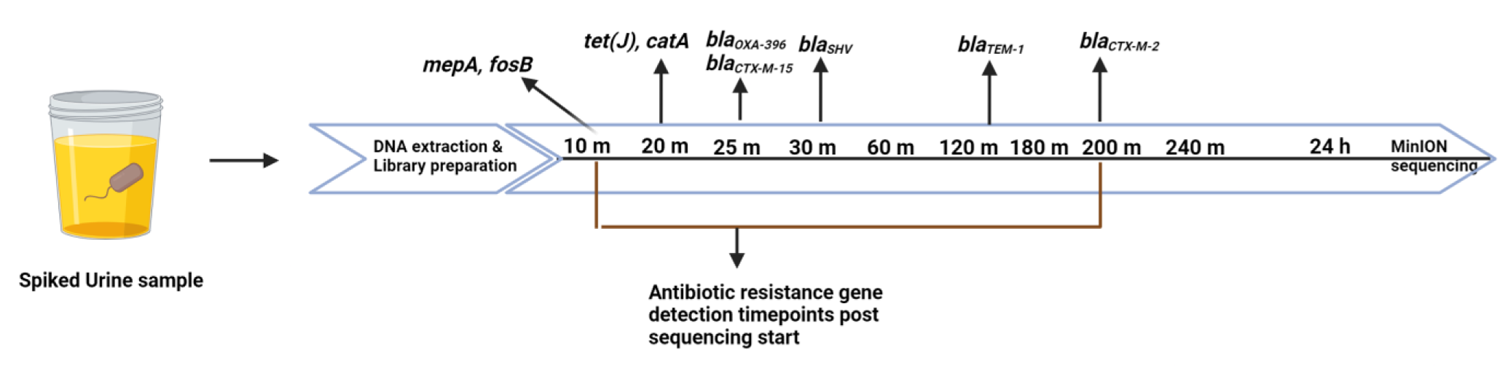
Detection of reference AMR genes over time for samples spiked with 10^5^ CFU/mL. Most of the genes were detected within 30 minutes of sequencing except blaTEM-1 and blaCTX-M-2, which took 140 and 200 minutes, respectively. BlaCTX-M-15 was detected after 13 hrs of sequencing from the sample spiked with 10^3^ CFU/mL.

In the case of *E. coli* NCTC 13441, the target AMR gene *bla_CTX-M-15_* was detected in samples spiked with both 10^3^ and 10^5^ CFU/mL concentrations of the bacteria. In samples spiked at 10^5^ CFU/mL, it was detected in 25 minutes, while the detection time frame was extended to 13 hours in samples spiked at 10^3^ CFU/mL. Moreover, other variants belonging to the *bla_CTX-M-15_* group were also detected in samples at both spike concentrations (2 in 10^3^ CFU/mL and 19 in 10^5^ CFU/mL).

For *E. coli* A2-39, the target AMR gene, *bla_CTX-M-2_*, was detected within 200 minutes, while an additional variant of the gene, *bla_CTX-M-35_*, was also detected. Similarly, for *E. coli* 155, the target AMR gene *bla_TEM-1_ and* one other variant of the same gene, *bla_TEM-234,_* were detected within 137 minutes. Seven variants of the target gene *bla_SHV_* were detected from samples spiked with 10^5^ CFU/mL of *K. pneumoniae*, but the specific variant of interest, *bla_SHV-11,_* was not identified. In the case of *P. mirabilis,* both the reference genes, *catA* and *tet(J),* were detected within 20 minutes of sequencing. For *P. aeruginosa*, the target gene *bla_oxa-356_* was detected within 25 minutes, and another of its variants, *bla_OXA-1_*. For *S. aureus* at 10^5^ CFU/mL, *fosB* and *mepA* genes were detected within 10 minutes, along with two other variants of the *mep* gene (*mepB* and *mepR*). Finally, no target ARGs were identified for samples spiked with INN 125 & *E. faecalis* CCUG 9997 in samples at either of the two tested spike concentrations.

### 3.5 Direct Nanopore Sequencing using Flongle flow cells

Five mL urine was individually spiked with three test strains, INN 125, *E. faecalis* CCUG9997 & *P. aeruginosa* CCUG17619 at the concentrations of 10^3^, 10^4^ & 10^5^ CFU/mL, and DNA samples obtained from direct DNA extraction were sequenced on Flongle flow cells. The raw data was base called in real-time using the integrated base caller Guppy, within the MinKNOW software. All the reads were considered for determining the pathogen ID and antibiotic resistance gene detection. A summary of the results is presented in Table III Table II & Figure V. All three pathogens were identified only in samples from the 10^5^ CFU/mL spike concentration, while none of the target antibiotic resistance genes were identified in samples at any of the three spiked concentrations.

**Figure V:**
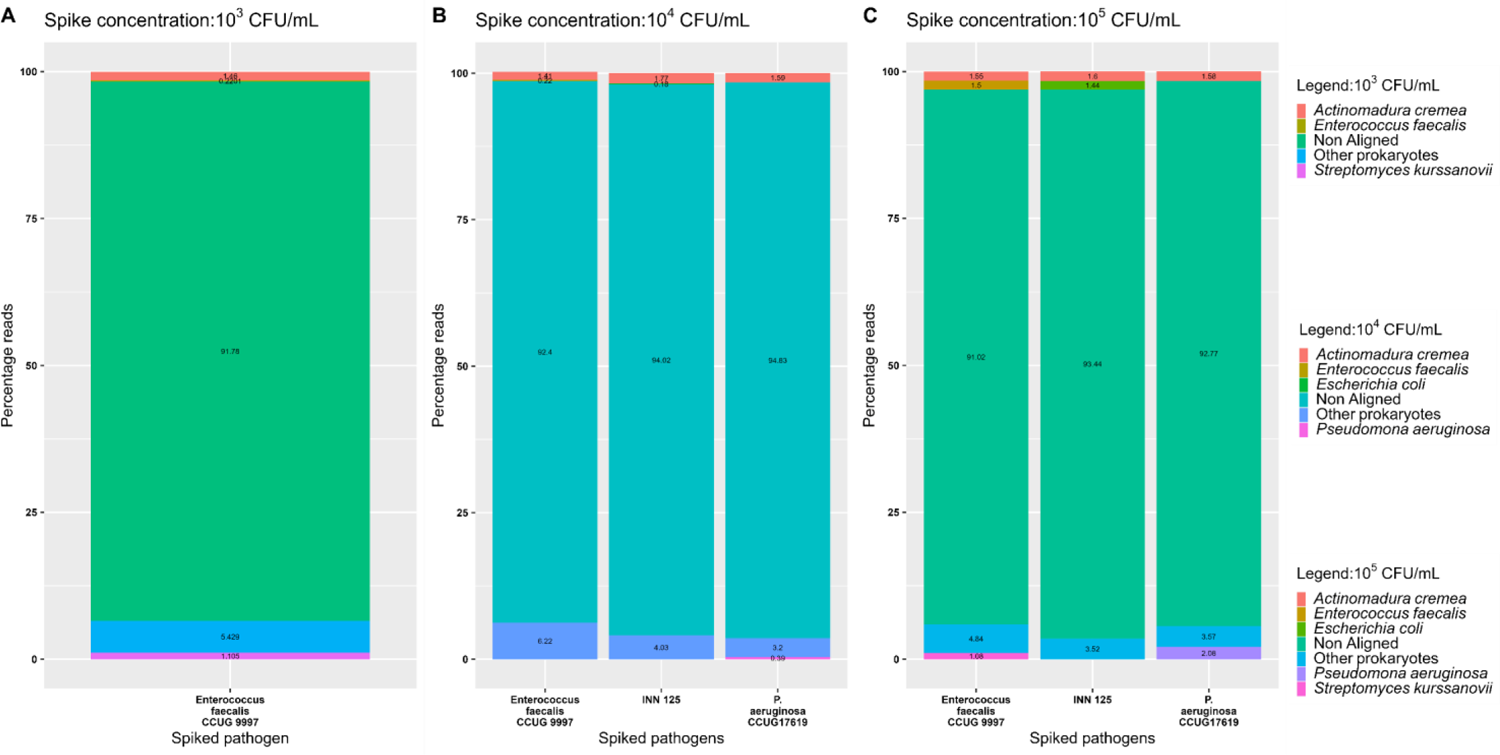
The figure denotes the distribution of reads in the sequence data generated by nanopore sequencing of DNA purified from spiked urine samples on Flongle flow cells. The subfigure A, B & C represent the samples spiked at 10^3^, 10^4^ & 10^5^ CFU/mL respectively. The represented percentage reads are based on a BLAST search against the RefProk database (prokaryotic sequence data only). The bacterial species (except the spiked pathogen) that were identified and had less than 1% of the total assigned reads were grouped and are represented as “other prokaryotes”.

**Table III:** Summary of the Pathogen Identification and AMR gene detection by Flongle sequencing from three different concentrations of 5 mL spiked urine samples. The legend for the figure is as follows: green tick: identification, red cross: negative identification & NA: no/unquantifiable DNA was isolated, and samples were excluded from the sequencing.

| Spiked pathogens | CFU/mL | DNA (ng/μl) | Pathogen Id | ARG |
| --- | --- | --- | --- | --- |
| *E. faecalis* CCUG9997 | 10<sup>3</sup> | 0.100 | × | × |
|  | 10<sup>4</sup> | 0.120 | × | × |
|  | 10<sup>5</sup> | 0.160 | ✓ | × |
| *P. aeruginosa* CCUG17619 | 10<sup>3</sup> | NA | - | - |
|  | 10<sup>4</sup> | 0.300 | × | × |
|  | 10<sup>5</sup> | 0.304 | ✓ | × |
| INN 125 | 10<sup>3</sup> | NA | - | - |
|  | 10<sup>4</sup> | 0.270 | × | × |
|  | 10<sup>5</sup> | 0.310 | ✓ | × |

The criteria for pathogen identification are the same as those used for MinION data analyses. However, the samples spiked with 10^3^ CFU/mL of *P. aeruginosa* CCUG17619 & INN 125 didn’t yield any DNA after the extraction steps. The results also indicate that 1-3% of reads identified as *Actinomadura cremea* were observed in all samples (except the 10^3^ CFU/mL of *P. aeruginosa* CCUG17619 & INN 125) from all three spike concentrations. The remaining identified bacterial species with less than 1% assigned reads were collectively presented as other prokaryotes.

All the samples exhibit a relatively higher number of non-aligned reads across all three concentrations. At 10^5^ CFU/mL of spike concentration, 1.44% of *E. coli*, 1.5% of *E. faecalis,* and 2.08% of *P. aeruginosa* reads were obtained from their respective spike samples. In all three cases, the highest number of prokaryotic reads were from the spiked pathogens, and pathogen identification was possible at 10^5^ CFU/mL of spike concentration. Comparatively, at 10^4^ CFU/mL of spike concentration, only 0.18% of *E. coli*, 0.22% of *E. faecalis,* and 0.39% of *P. aeruginosa* reads were obtained from their respective spike samples, and the percentage of reads remained the same for *E. faecalis* at 10^3^ CFU/mL spike concentration. Hence, identifying the spiked pathogens was impossible at 10^3^ & 10^4^ CFU/mL concentrations. The identification of antibiotic-resistance genes using the ABRicate tool didn’t yield any positive results, and no target genes were identified at any of the spike concentrations.

### 3.6 Genome coverage analysis at different concentrations of spiking

The raw sequencing reads obtained per bin were aligned with the reference genome of the respective spiked species to observe the genome coverage over time (Figure VI). Among the species spiked with 10^5^ CFU/mL, the genome coverage was > 90 % in five spiked samples corresponding to E. coli NCTC 13441, *K. pneumoniae* CCUG 255T*, P. mirabilis* CCUG 26767T*, P. aeruginosa* CCUG 17619 *and S. aureus* NCTC 8325 (Figure 6. B). For the samples spiked with 10^3^ CFU/mL of the four gram-negative species, *E. coli, K. pneumoniae, P. mirabilis,* and *P. aeruginosa* was around 10%, while the coverage of gram-positive *E. faecalis* and *S. aureus* was less than 1%. *P. aeruginosa* had a coverage similar to the other three gram-negative strains however, it was not detected from the 10^3^ CFU/mL. The genome coverage of three in-house strains at 10^5^ CFU/mL was 25.7 % for INN 125, 37.6 % for INN A2-39, and around 7 % for INN 155. Similarly, the genome coverage at 10^3^ CFU/mL was below 5% for all of them (Supplementary Figure V).

**Figure VI:**
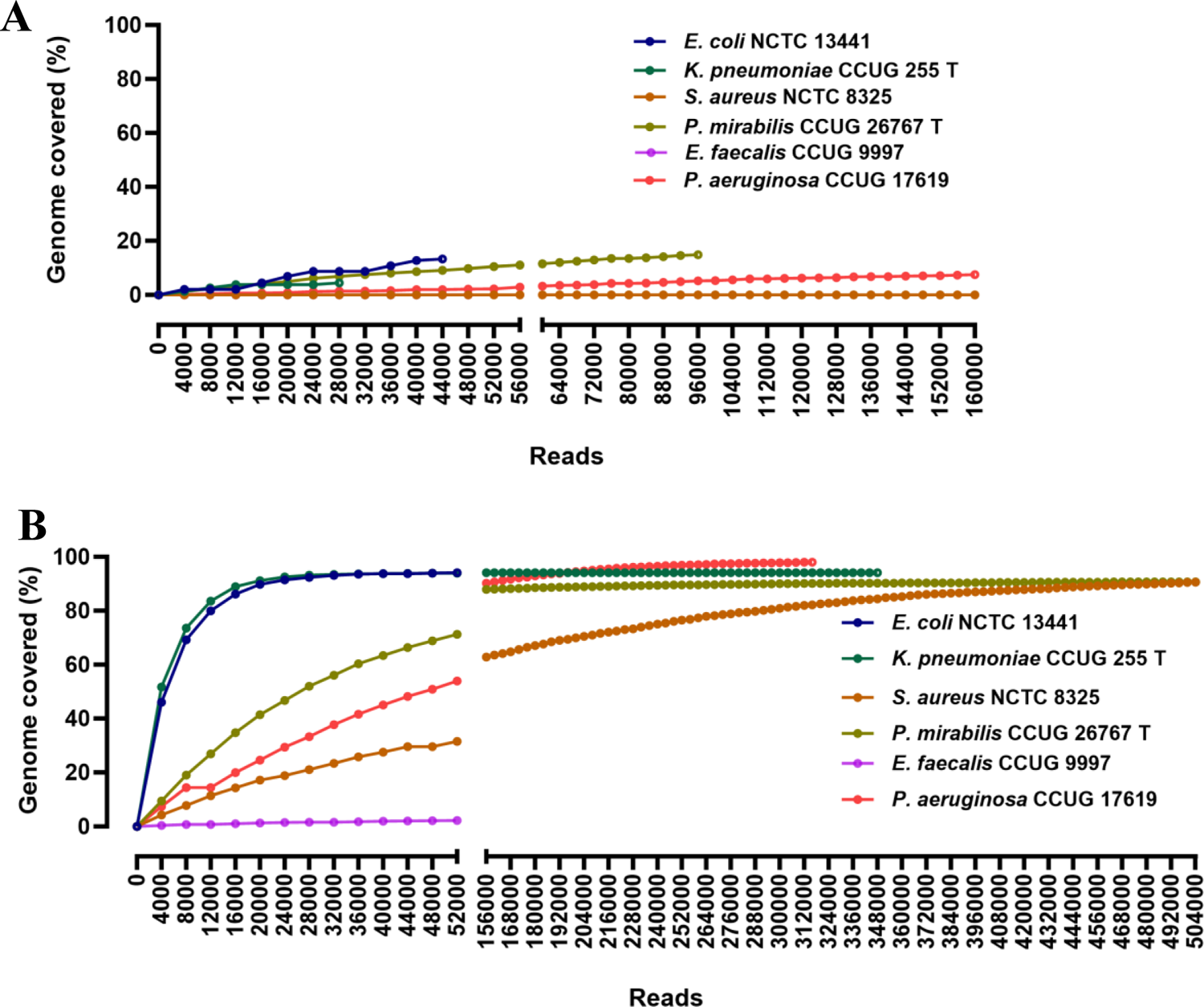
Genome coverage of the target species over time. The subfigure 6. A represents the pathogen coverage obtained for the samples spiked with 10^3^ CFU/mL concentration, while the subfigure 6.B. is the pathogen coverage from the samples spiked with a pathogen concentration of 10^5^ CFU/mL.

## 4 Discussion

Rapid and accurate diagnosis of pathogens and their resistance profiles is critical for administering effective antibiotics. This proof-of-concept study aimed to establish a rapid, culture-and-amplification-free method of detecting pathogens and their corresponding antibiotic-resistance genes through Nanopore sequencing from spiked urine samples. An OD-CFU-based methodology was developed for mimicking the clinical urine samples. DNA isolation optimization in which three commercial DNA extraction kits, namely QIAamp BiOstic Bacteremia DNA Kit, PowerFood Microbial Kit and DNeasy Blood & Tissue Kit identified the latter as an optimal kit when the factors of DNA yield, protocol length and cost per sample were considered. Extracted DNA was sequenced on MinION and Flongle platforms. Except for INN 125 and *E. faecalis* CCUG9997, all the spike strains utilized, and their encoded reference antibiotic resistance genes were identified in samples spiked at 10^5^ CFU/mL on MinION flow cells, while only pathogen identification was possible in samples at the same spike concentration using Flongle sequencing. Meanwhile, for the 10^3^ CFU/mL spiked samples, identifying pathogens and antibiotic resistance genes was possible only for *E. coli NCTC 13441* through MinION sequencing. In contrast, no pathogens and antibiotic resistance genes were identified through Flongle sequencing.

### 4.1 A combination of bacterial optical densities, colony forming units (CFU), and plate count methods for spiking healthy urine to mimic clinical samples

The present study uses a correlation between the optical densities and the cell concentration for mimicking the UTI samples using fresh urine samples at two different concentrations of 10^3^ & 10^5^ CFU/mL. In most cases, UTI is considered microbiologically confirmed when the bacterial count in urine culture reaches 10^5^ CFU/mL. However, lower concentrations, such as 10^3^ and even 10^2^ CFU/mL might still be clinically relevant, particularly in cases of cystitis in children (36).

The optical density didn’t always correlate with the bacterial CFU count. The discrepancy is also reported in different interlaboratory studies (37–39), which have highlighted the poor relationship between OD measurements and CFU counts in terms of reproducibility and reliability. On the other hand, some studies have concluded that OD measurements directly correlate with cell concentration and can be reliable and reproducible (40). However, these studies accepted error rates higher than 50% to arrive at this conclusion. The spiked suspension was always cultured to overcome the bias in the correlation between OD and CFU count. This resulted in the determination of the exact bacterial concentration of the prepared spike solution. While the colony count method effectively verifies the bacterial titer in solution, one needs to remember that this method only allows the enumeration of viable bacteria. Hence, a fresh overnight-grown culture must be used to minimize experimental errors.

### 4.2 Direct DNA extraction from spiked urine samples

DNA extraction is a crucial step for sequencing-based analysis, and the present study evaluated three different commercial kits: BiOstic, Power Food, and Blood and Tissue. The kits were compared in terms of DNA yield, protocol length, and cost per sample. Among gram-negative bacteria spiked samples, the concentration of DNA isolated by the DNeasy Blood and Tissue kit from *E. coli* and *K. pneumoniae* spiked samples was comparatively higher in all three tested concentrations of CFU/mL. Similarly, among *S. aureus* (gram-positive) samples, the concentration of DNA extracted using a Biostic kit from 10^3^ and 10^4^ CFU/mL spike samples (0.75 ng/µl and 0.64 ng/µl, respectively) was slightly higher than that of those for DNeasy Blood and Tissue kit (0.57 ng/µl and 0.58 ng/µl, respectively). Although the DNeasy Blood & Tissue kit has lower per microlitre concentrations in the *S. aureus* samples and in the 10^5^ CFU/mL spiked samples of *K. pneumoniae*, the DNA yield obtained through this kit is higher than the Biostic kit as the elution volume of the former (200 µl) is four times higher than that of the latter (50 µl). Hence, sufficient DNA was extracted from the spiked samples for the sequencing steps within 35 minutes. Previous studies have also reported that the Blood and Tissue kit is best for DNA isolation from artificial human urine samples and canine urine (41,42). Karstens et al. (43) also reported higher DNA yield from human urine by the Blood and Tissue kit, although DNA produced by both BiOstic and Blood and Tissue kits performed well in downstream analysis. Moreover, the blood and tissue kit costs only 3.94 € per sample, which is lower than the 4.29 € and 4.64 € per sample costs for Power Food and BiOstic kits.

### 4.3 Identification of pathogens and antibiotic resistance genes by MinION and Flongle sequencing

All the pathogenic strains used in this study are common uropathogens and were selected based on their occurrence in complicated urinary tract infections. These spiked pathogens apart from INN 125, INN A2-39 *&* INN 155 are standard strains, and the reference assemblies for these strains are available in public databases. Meanwhile, the three stated clinical in-house strains (INN 125, INN A2-39 *&* INN 155) were sequenced on Illumina and Nanopore platforms, and hybrid assemblies were created for these strains. All the pathogens were accurately identified in samples with 10^5^ CFU/mL spike concentration, while only three species were identified at 10^3^ CFU/mL spike concentration. The pathogen identification in both spike concentrations was obtained within 10 minutes of sequencing start, from the first 4000 reads.

Numerous studies have reported on the use of MinION-based metagenomic sequencing for pathogens and antibiotic resistance gene identification from biological fluids such as blood (44), cerebrospinal fluid (45) and urine samples (18,19,46,47). Schmidt et al. (19) reported the identification of pathogens by MinION sequencing directly from the urines with a turnaround time of 4 hours. However, the samples used for the analysis were heavily infected clinical samples with CFU counts >10^7^ CFU/mL.f

Our results show that pathogen identification becomes challenging in samples with 10^3^ CFU/mL spikes, indicating that this is probably the limit of detection for the current methodology. This observation resonates with the existing literature where the nanopore sequencing-based assays have a limit of detection between 10^4^ to 10^2^ CFU/mL (48,49). Authors Zhang et al. reported the identification of the pathogen from urine samples spiked with ≥10^4^ CFU/mL of *E. coli* and *Enterococcus faecium* within 2 hours of sequencing, while only *E. faecium* was detected from 10^3^ CFU/mL spiked samples. Similarly, authors Deng et al. determined the limit of detection to be 10^3^ CFU/mL for *E. coli* in spiked sputum samples. However, they used amplicon sequencing, a targeted sequencing method that amplifies specific DNA regions using PCR before sequencing (51). Similar to MinION-based sequencing, a subset of the strains used in the study were spiked at three concentrations of 10^3^, 10^4^ & 10^5^ CFU/mL. Flongle flow cells are the cost-effective version of the MinION flow cell, hence facilitating the reduction of sequencing costs per run. The obtained results for the pathogen identification at 10^5^ CFU/mL complement the MinION-based results at the same concentration but failed to detect the relevant reference antibiotic resistance genes.

### 4.4 Diagnosing clinically relevant antibiotic-resistance genes through metagenomic sequencing depends on bacterial sequence coverage

The strains used in the current study are all well characterized; hence, the ground truth about the presence of antibiotic-resistance genes is known. In the current study, most of the reference antibiotic resistance genes were identified within half an hour of the sequencing start on the MinION-based sequencing. At a 10^5^ CFU/mL spike concentration, seven of the total nine reference genes were detected, while at 10^3^ CFU/mL, detection was possible in only one sample after 13 hours of sequencing. Meanwhile, no antibiotic-resistance genes were detected from any Flongle sequencing runs. The ability to identify antibiotic resistance genes from the metagenomic data suggests that enough sequencing depth is achieved for sufficient genome coverage of the target pathogen, as a high level of coverage is required for robust detection.

The MinION sequencing results indicate that a genome coverage of over 90% was achieved among *E. coli* NCTC 13441, *K. pneumoniae CCUG225T, P. mirabilis CCUG2676T, P. aeruginosa CCUG17619* and *S. aureus NCTC8325* species at the concentration of 10^5^ CFU/mL. All the corresponding antibiotic resistance genes (*bla_CTX-M-15_, bla_CTX-M-2_, bla_SHV_, tet(J), mepA, bla_OXA-396_, fosB*) present in these spiked strains were identified between 30 to 200 minutes of sequencing start. The unidentified resistance genes *bla_TEM-1_* and *tet(M)* present in strains INN 125 and *E. faecalis* were missed due to lower percentage coverage. Similar results were observed for 10^3^ CFU/mL, as the coverage is less than 10% in all the samples. The same can be concluded for the Flongle results as the coverage is far poorer compared to MinION-based results, which has also been observed in our previous study (52).

In summary, the current study established an OD-CFU-based method for spiking healthy urine samples to mimic clinical samples and establishing a pipeline for direct DNA extraction from these spiked samples. The performances of BiOstic, PowerFood, and Blood & tissue kits were compared for DNA extraction. The study determined that the performance of the blood and tissue kit was better than the other kits based on the DNA yield, length of extraction, and cost per sample. All the spiked pathogens at 10^5^ CFU/mL concentrations were identified directly through MinION sequencing within 1.5 hours of sample processing. The first read file with 4000 reads generated in 10 minutes of sequencing was enough for identification, while three hours of sequencing were required for antibiotic resistance gene detection. Hence, both the pathogen identification and ARG detection were possible within 5 hours of sample processing through MinION sequencing. Additionally, pathogen identification for *E. coli, K. pneumoniae,* and *P. mirabilis* was possible at 10^3^ CFU/mL using MinION sequencing. The coverage for these samples was less than 15%, indicating that most of the isolated and sequenced DNA probably belonged to the host. Moreover, extracting sufficient quantities of pathogenic DNA at such low CFU becomes challenging as they are present at lower abundance levels (53).

Additionally, this study had some limitations. Firstly, the urine volume used for DNA extraction is 30 ml. Clinically, obtaining such a large volume of urine from critically ill patients with reduced kidney function might be challenging. Secondly, we have also observed difficulties extracting spiked *E. faecalis* samples as the DNA yield was insufficient for downstream analysis.

In conclusion, this proof-of-concept study shows that detecting the pathogen and antibiotic resistance genes was possible through direct DNA extraction and sequencing on MinION flow cells with a turn-around time of five hours. This method can significantly reduce the conventional turnaround time from approximately 48 hours to 5 hours, aiding in the administration of tailored and more effective antibiotics, thereby increasing the chance of patient survival, and preventing the spread of AMR.

## Supporting information

Supplementary Figure III

supplementary table III

## Reference

1. Newlands AF, Roberts L, Maxwell K, Kramer M, Price JL, Finlay KA. The Recurrent Urinary Tract Infection Symptom Scale: Development and validation of a patient-reported outcome measure. BJUI Compass. 2023;4(3):285– 97.

2. Mortazavi-Tabatabaei SAR, Ghaderkhani J, Nazari A, Sayehmiri K, Sayehmiri F, Pakzad I. Pattern of Antibacterial Resistance in Urinary Tract Infections: A Systematic Review and Meta-analysis. Int J Prev Med. 2019;10:169.

3. Yang X, Chen H, Zheng Y, Qu S, Wang H, Yi F. Disease burden and long-term trends of urinary tract infections: A worldwide report. Frontiers in Public Health. 2022;10:888205.

4. Zeng Z, Zhan J, Zhang K, Chen H, Cheng S. Global, Regional, and National Burden of Urinary Tract Infections from 1990-2019: An Analysis of the Global Burden of Disease Study [Internet]. 2021. Available from: 10.21203/rs.3.rs-829349/v1

5. Sabih A, Leslie SW. Complicated Urinary Tract Infections. In: StatPearls [Internet]. Treasure Island (FL): StatPearls Publishing; 2024 [cited 2024 Feb 26]. Available from: http://www.ncbi.nlm.nih.gov/books/NBK436013/

6. Guliciuc M, Maier AC, Maier IM, Kraft A, Cucuruzac RR, Marinescu M, et al. The Urosepsis-A Literature Review. Medicina (Kaunas). 2021 Aug 25;57(9):872.

7. Ryan J, O’Neill E, McLornan L. Urosepsis and the urologist! Current Urology. 2021;15(1):39–44.

8. Foxman B. Urinary tract infection syndromes: occurrence, recurrence, bacteriology, risk factors, and disease burden. Infect Dis Clin North Am. 2014 Mar;28(1):1–13.

9. Medina-Polo J, Naber KG, Bjerklund Johansen TE. Healthcare-associated urinary tract infections in urology. GMS Infectious Diseases; 9:Doc05 [Internet]. 2021 Aug 30 [cited 2023 May 29]; Available from: https://www.egms.de/en/journals/id/2021-9/id000074.shtml

10. Dias V. Candida species in the urinary tract: is it a fungal infection or not? Future Microbiology. 2020 Jan;15(2):81– 3.

11. Bilsen MP, Jongeneel RMH, Schneeberger C, Platteel TN, van Nieuwkoop C, Mody L, et al. Definitions of Urinary Tract Infection in Current Research: A Systematic Review. Open Forum Infect Dis. 2023 Jun 27;10(7):ofad332.

12. Li W, Sun E, Wang Y, Pan H, Zhang Y, Li Y, et al. Rapid Identification and Antimicrobial Susceptibility Testing for Urinary Tract Pathogens by Direct Analysis of Urine Samples Using a MALDI-TOF MS-Based Combined Protocol. Front Microbiol. 2019 Jun 5;10:1182.

13. Cheng K, Chui H, Domish L, Hernandez D, Wang G. Recent development of mass spectrometry and proteomics applications in identification and typing of bacteria. Proteomics Clin Appl. 2016 Apr;10(4):346–57.

14. Shin DJ, Andini N, Hsieh K, Yang S, Wang TH. Emerging Analytical Techniques for Rapid Pathogen Identification and Susceptibility Testing. Annual Review of Analytical Chemistry. 2019;12(1):41–67.

15. Haider A, Ringer M, Kotroczó Z, Mohácsi-Farkas C, Kocsis T. The Current Level of MALDI-TOF MS Applications in the Detection of Microorganisms: A Short Review of Benefits and Limitations. Microbiology Research. 2023 Mar;14(1):80–90.

16. Wang Y, Chen T, Zhang S, Zhang L, Li Q, Lv Q, et al. Clinical evaluation of metagenomic next-generation sequencing in unbiased pathogen diagnosis of urinary tract infection. Journal of Translational Medicine [Internet]. 2023 [cited 2024 Feb 15];21. Available from: https://www.ncbi.nlm.nih.gov/pmc/articles/PMC10612365/

17. Grumaz C, Hoffmann A, Vainshtein Y, Kopp M, Grumaz S, Stevens P, et al. Rapid Next-Generation Sequencing– Based Diagnostics of Bacteremia in Septic Patients. The Journal of Molecular Diagnostics. 2020 Mar 1;22(3):405–18.

18. Zhang L, Huang W, Zhang S, Li Q, Wang Y, Chen T, et al. Rapid Detection of Bacterial Pathogens and Antimicrobial Resistance Genes in Clinical Urine Samples With Urinary Tract Infection by Metagenomic Nanopore Sequencing. Front Microbiol. 2022 May 17;13:858777.

19. Schmidt K, Mwaigwisya S, Crossman LC, Doumith M, Munroe D, Pires C, et al. Identification of bacterial pathogens and antimicrobial resistance directly from clinical urines by nanopore-based metagenomic sequencing. J Antimicrob Chemother. 2017 Jan;72(1):104–14.

20. Liu M, Yang S, Wu S, Chen L, Li S, Li Z, et al. Detection of pathogens and antimicrobial resistance genes directly from urine samples in patients suspected of urinary tract infection by metagenomics nanopore sequencing: A large-scale multi-centre study. Clinical and Translational Medicine. 2023;13(4):824.

21. Vendrell JA, Henry S, Cabello-Aguilar S, Heckendorn E, Godreuil S, Solassol J. Determination of the Optimal Bacterial DNA Extraction Method to Explore the Urinary Microbiota. International Journal of Molecular Sciences. 2022 Jan;23(3):1336.

22. Fiedorová K, Radvanský M, Němcová E, Grombiříková H, Bosák J, Černochová M, et al. The Impact of DNA Extraction Methods on Stool Bacterial and Fungal Microbiota Community Recovery. Frontiers in Microbiology [Internet]. 2019 [cited 2024 Feb 15];10. Available from: https://www.frontiersin.org/journals/microbiology/articles/10.3389/fmicb.2019.00821

23. Ackerman AL, Anger JT, Khalique MU, Ackerman JE, Tang J, Kim J, et al. Optimization of DNA extraction from human urinary samples for mycobiome community profiling. PLOS ONE. 2019 Apr 25;14(4):e0210306.

24. Karstens L, Siddiqui NY, Zaza T, Barstad A, Amundsen CL, Sysoeva TA. Benchmarking DNA isolation kits used in analyses of the urinary microbiome. Scientific Reports. 2021;11(1):6186.

25. Zhang L, Chen T, Wang Y, Zhang S, Lv Q, Kong D, et al. Comparison Analysis of Different DNA Extraction Methods on Suitability for Long-Read Metagenomic Nanopore Sequencing. Frontiers in Cellular and Infection Microbiology [Internet]. 2022 [cited 2023 Dec 8];12. Available from: https://www.frontiersin.org/articles/10.3389/fcimb.2022.919903

26. Ali J, Johansen W, Ahmad R. Short turnaround time of seven to nine hours from sample collection until informed decision for sepsis treatment using nanopore sequencing. Scientific Reports. 2024;

27. Cheesbrough M. District Laboratory Practice in Tropical Countries [Internet]. 2nd ed. Cambridge University Press; 2006. Available from: 10.1017/CBO9780511543470

28. Munch MM, Chambers LC, Manhart LE, Domogala D, Lopez A, Fredricks DN, et al. Optimizing bacterial DNA extraction in urine. PLoS One. 2019 Sep 24;14(9):e0222962.

29. Camacho C, Coulouris G, Avagyan V, Ma N, Papadopoulos J, Bealer K, et al. BLAST+: architecture and applications. BMC Bioinformatics. 2009 Dec 15;10(1):421.

30. Avershina E, Khezri A, Ahmad R. Clinical Diagnostics of Bacterial Infections and Their Resistance to Antibiotics— Current State and Whole Genome Sequencing Implementation Perspectives. Antibiotics. 2023 Apr;12(4):781.

31. Seemann T. Abricate [Internet]. 2017 [cited 2024 Feb 26]. Available from: https://github.com/tseemann/abricate

32. Jia B, Raphenya AR, Alcock B, Waglechner N, Guo P, Tsang KK, et al. CARD 2017: Expansion and model-centric curation of the comprehensive antibiotic resistance database. Nucleic Acids Research. 2017;45(D1):566–73.

33. Zankari E, Hasman H, Cosentino S, Vestergaard M, Rasmussen S, Lund O, et al. Identification of acquired antimicrobial resistance genes. Journal of Antimicrobial Chemotherapy. 2012;67(11):2640–4.

34. Doster E, Lakin SM, Dean CJ, Wolfe C, Young JG, Boucher C, et al. MEGARes 2.0: A database for classification of antimicrobial drug, biocide and metal resistance determinants in metagenomic sequence data. Nucleic Acids Research. 2020;48(D1):561–9.

35. Feldgarden M, Brover V, Haft DH, Prasad AB, Slotta DJ, Tolstoy I, et al. Validating the AMRFinder Tool and Resistance Gene Database by Using Antimicrobial Resistance Genotype-Phenotype Correlations in a Collection of Isolates. Antimicrobial Agents and Chemotherapy. 2019;63(11):00483–19.

36. De Cueto M, Aliaga L, Alós JI, Canut A, Los-Arcos I, Martínez JA, et al. Executive summary of the diagnosis and treatment of urinary tract infection: Guidelines of the Spanish Society of Clinical Microbiology and Infectious Diseases (SEIMC). Enfermedades Infecciosas y Microbiología Clínica. 2017 May;35(5):314–20.

37. Beal J, Farny NG, Haddock-Angelli T, Selvarajah V, Baldwin GS, Buckley-Taylor R, et al. Robust estimation of bacterial cell count from optical density. Communications Biology. 2020;3(1):512.

38. McBirney SE, Trinh K, Wong-Beringer A, Armani AM. Wavelength-normalized spectroscopic analysis of Staphylococcus aureus and Pseudomonas aeruginosa growth rates. Biomedical Optics Express. 2016;7(10):4034.

39. Jarvis B, Hedges AJ, Corry JEL. Assessment of measurement uncertainty for quantitative methods of analysis: Comparative assessment of the precision (uncertainty) of bacterial colony counts. International Journal of Food Microbiology. 2007;116(1):44–51.

40. Biesta-Peters EG, Reij MW, Joosten H, Gorris LGM, Zwietering MH. Comparison of Two Optical-Density-Based Methods and a Plate Count Method for Estimation of Growth Parameters of Bacillus cereus. Applied and Environmental Microbiology. 2010;76(5):1399–405.

41. Mrofchak R, Madden C, Evans MV, Hale VL. Evaluating extraction methods to study canine urine microbiota. Gyarmati P, editor. PLoS ONE. 2021 Jul 9;16(7):e0253989.

42. Vendrell JA, Henry S, Cabello-Aguilar S, Heckendorn E, Godreuil S, Solassol J. Determination of the Optimal Bacterial DNA Extraction Method to Explore the Urinary Microbiota. IJMS. 2022 Jan 25;23(3):1336.

43. Karstens L, Siddiqui NY, Zaza T, Barstad A, Amundsen CL, Sysoeva TA. Benchmarking DNA isolation kits used in analyses of the urinary microbiome. Sci Rep. 2021 Mar 17;11(1):6186.

44. Ren D, Ren C, Yao R, Zhang L, Liang X, Li G, et al. The microbiological diagnostic performance of metagenomic next-generation sequencing in patients with sepsis. BMC Infect Dis. 2021 Dec; 21(1):1257.

45. Wilson MR, Naccache SN, Samayoa E, Biagtan M, Bashir H, Yu G, et al. Actionable diagnosis of neuroleptospirosis by next-generation sequencing. N Engl J Med. 2014 Jun 19;370(25):2408–17.

46. Liu M, Yang S, Wu S, Chen L, Li S, Li Z, et al. Detection of pathogens and antimicrobial resistance genes directly from urine samples in patients suspected of urinary tract infection by metagenomics nanopore sequencing: A large-scale multi-centre study. Clinical & Translational Med. 2023 Apr;13(4):e824.

47. Deng Q, Cao Y, Wan X, Wang B, Sun A, Wang H, et al. Nanopore-based metagenomic sequencing for the rapid and precise detection of pathogens among immunocompromised cancer patients with suspected infections. Frontiers in Cellular and Infection Microbiology. 2022;12:943859.

48. Player R, Verratti K, Staab A, Bradburne C, Grady S, Goodwin B, et al. Comparison of the performance of an amplicon sequencing assay based on Oxford Nanopore technology to real-time PCR assays for detecting bacterial biodefense pathogens. BMC Genomics. 2020 Feb 1):166.

49. Lewandowski K, Xu Y, Pullan ST, Lumley SF, Foster D, Sanderson N, et al. Metagenomic Nanopore Sequencing of Influenza Virus Direct from Clinical Respiratory Samples. J Clin Microbiol. 2019 Dec 23;58(1):e00963–19.

50. Zhang L, Huang W, Zhang S, Li Q, Wang Y, Chen T, et al. Rapid Detection of Bacterial Pathogens and Antimicrobial Resistance Genes in Clinical Urine Samples With Urinary Tract Infection by Metagenomic Nanopore Sequencing. Front Microbiol. 2022 May 17;13:858777.

51. Nygaard AB, Tunsjø HS, Meisal R, Charnock C. A preliminary study on the potential of Nanopore MinION and Illumina MiSeq 16S rRNA gene sequencing to characterize building-dust microbiomes. Sci Rep. 2020 Feb 21;10(1):3209.

52. Avershina E, Frye SA, Ali J, Taxt AM, Ahmad R. Ultrafast and Cost-Effective Pathogen Identification and Resistance Gene Detection in a Clinical Setting Using Nanopore Flongle Sequencing. Frontiers in Microbiology [Internet]. 2022 [cited 2023 Sep 11];13. Available from: https://www.frontiersin.org/articles/10.3389/fmicb.2022.822402

53. Knudsen BE, Bergmark L, Munk P, Lukjancenko O, Priemé A, Aarestrup FM, et al. Impact of Sample Type and DNA Isolation Procedure on Genomic Inference of Microbiome Composition. mSystems. 2016 Oct 18;1(5):10.1128/msystems.00095-16.

