## Supplementary Figure I for "Detection of pathogens and antimicrobial resistant genes from urine within 5 hours using Nanopore sequencing"

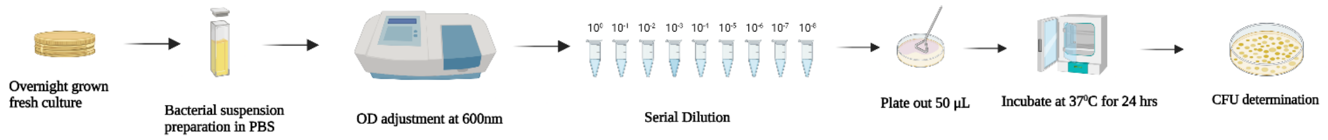


**Supplementary figure I:** A graphical overview of the different steps involved in the optical density and standard platting-based methods for inoculum optimisation.
