## Supplementary Figure II for "Detection of pathogens and antimicrobial resistant genes from urine within 5 hours using Nanopore sequencing"

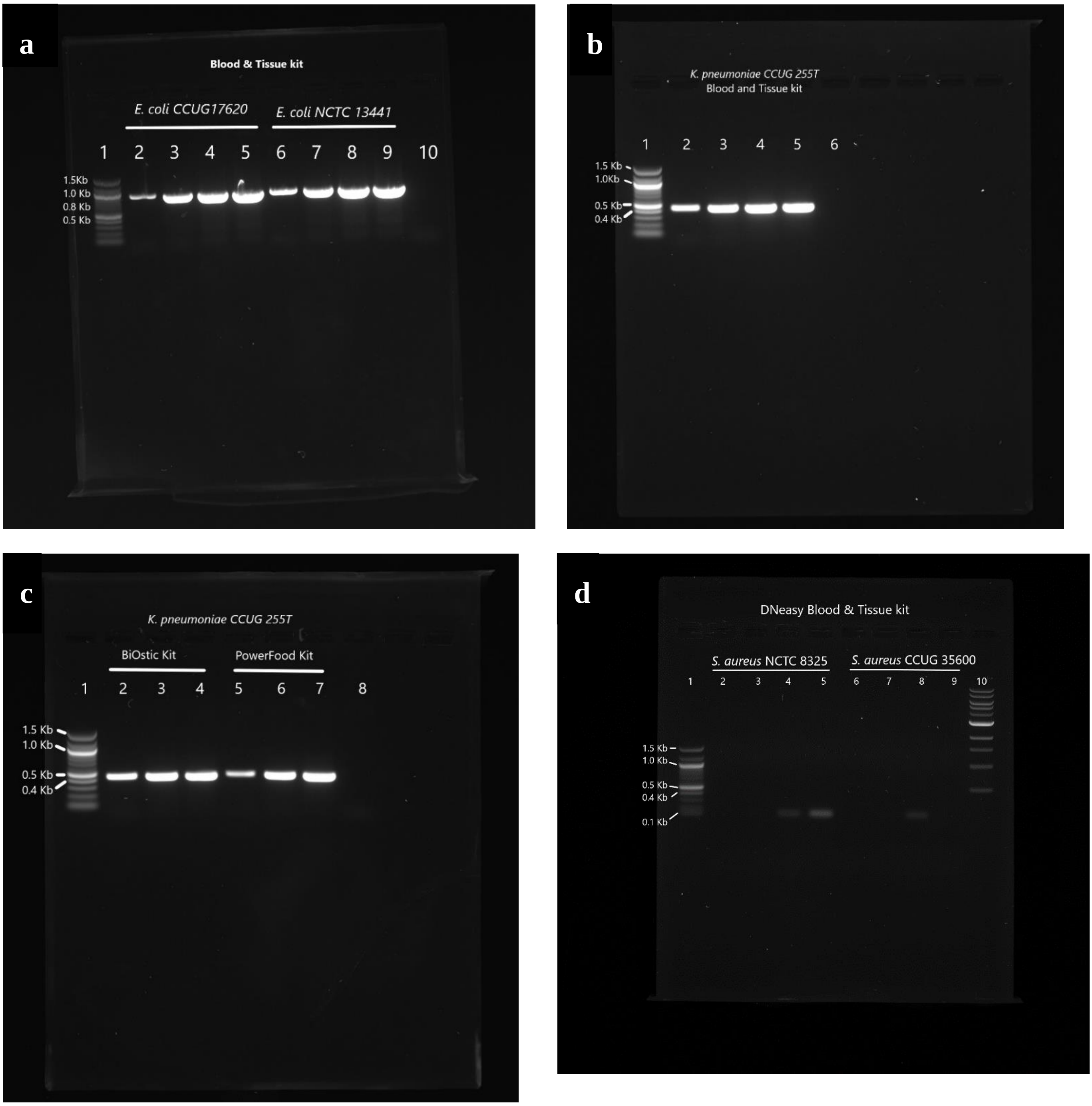
a
Blood & Tissue kit
E. coli CCUG17620
E. coli NCTC 13441
1 2 3 4 5 6 7 8 9 10
1.5Kb
1.0 Kb
0.8 Kb
0.5 Kb
b
K. pneumoniae CCUG 255T
Blood and Tissue kit
1 2 3 4 5 6
1.5 Kb
1.0Kb
0.5 Kb
0.4 Kb
c
K. pneumoniae CCUG 255T
BiOstic Kit
PowerFood Kit
1 2 3 4 5 6 7 8
1.5 Kb
1.0 Kb
0.5 Kb
0.4 Kb
d
DNeasy Blood & Tissue kit
S. aureus NCTC 8325
S. aureus CCUG 35600
1 2 3 4 5 6 7 8 9 10
1.5 Kb
1.0 Kb
0.5 Kb
0.4 Kb
0.1 Kb

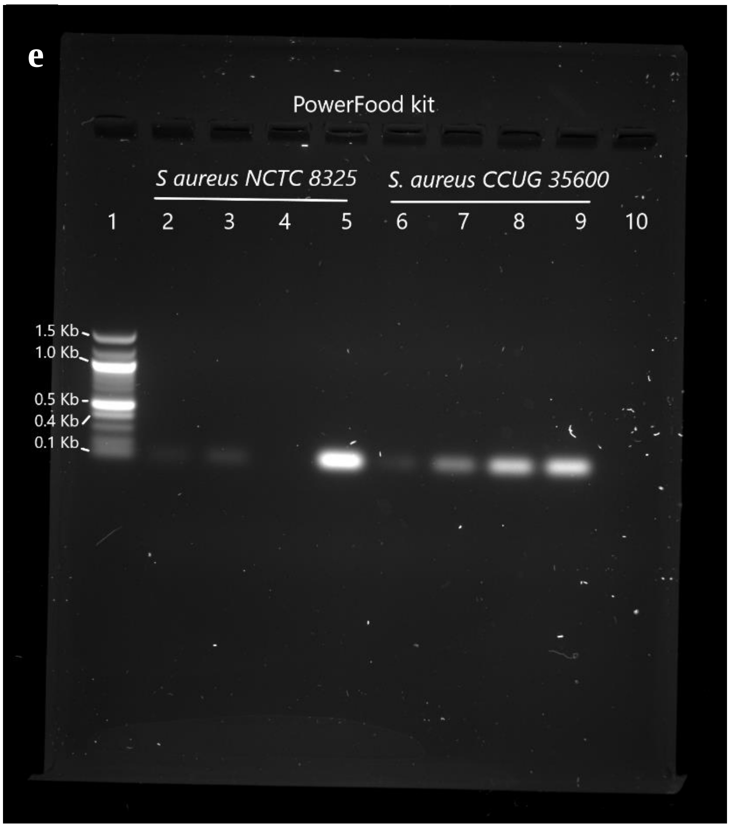


**Supplementary figure II:** PCR results for the DNA extracted from different kits at two different spike concentrations for *E. coli CCUG* 17620, *E. coli NCTC* 13441 and *K. pneumoniae* CCUG 255T. For the *E. coli* spiked samples, the amplification was performed using *UspA* gene (850 bp), while for *K. pneumoniae, Khe* (428 bp) housekeeping gene was used.

Lane description: **subfigure II. a.** 1: 100bp NEB DNA ladder, 3: $10^3$ CFU/mL, 5: $10^5$ CFU/mL, 7: $10^3$ CFU/mL, 9: $10^5$ CFU/mL, 10: Negative template control.

Lane description: **subfigure II. b**. 1: 100bp NEB DNA ladder, 2: $10^3$ CFU/mL, 4: $10^5$ CFU/mL, 6: Negative template control.

Lane description: **subfigure II. c**. 1: 100bp NEB DNA ladder, 2: $10^3$ CFU/mL, 4: $10^5$ CFU/mL, 5: $10^3$ CFU/mL, 7: $10^5$ CFU/mL, 8: Negative template control.

Lane description: **subfigure II. d.** 1: 100bp NEB DNA ladder, 2: $10^3$ CFU/mL, 4: $10^5$ CFU/mL, 6: $10^3$ CFU/mL, 8: $10^5$ CFU/mL, 10: 1kB DNA ladder.

Lane description: **subfigure II. e.** 1: 100bp NEB DNA ladder, 2: $10^3$ CFU/mL, 4: $10^5$ CFU/mL, 6: $10^3$ CFU/mL, 8: $10^5$ CFU/mL, 10: Negative template control.
