## Supplementary Figure III for "Detection of pathogens and antimicrobial resistant genes from urine within 5 hours using Nanopore sequencing"

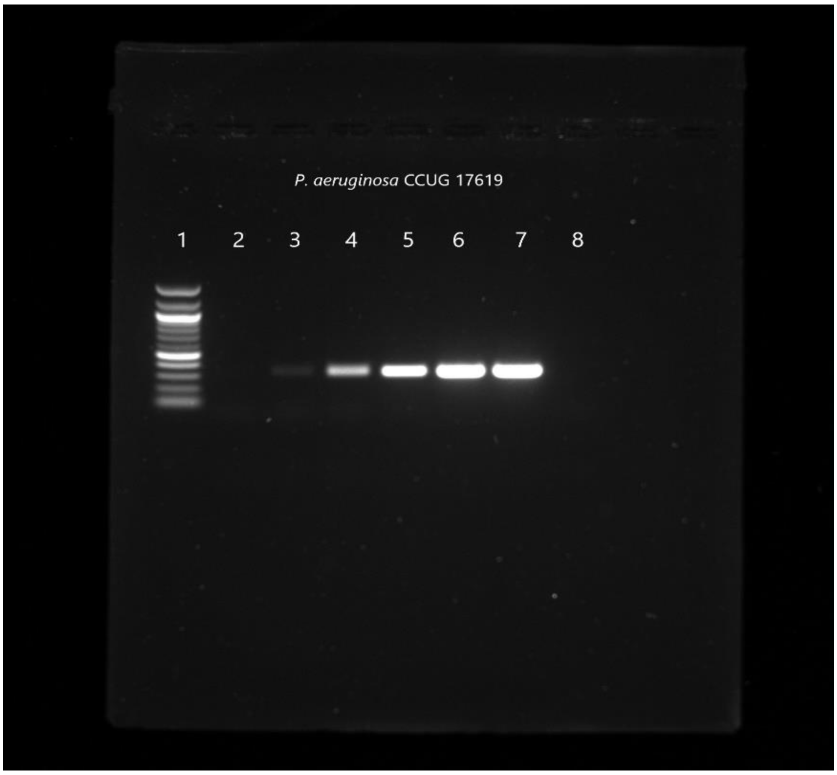


**Supplementary figure III:** PCR results for the DNA extracted from samples spiked with *P. aeruginosa* at concentration of $10^3$ & $10^5$ CFU/mL. Lane description: 1: 100bp NEB DNA ladder, 3: $10^3$ CFU/mL, 5: $10^5$ CFU/mL, 8: Negative template control.
