## Supplementary Figure IV for "Detection of pathogens and antimicrobial resistant genes from urine within 5 hours using Nanopore sequencing"

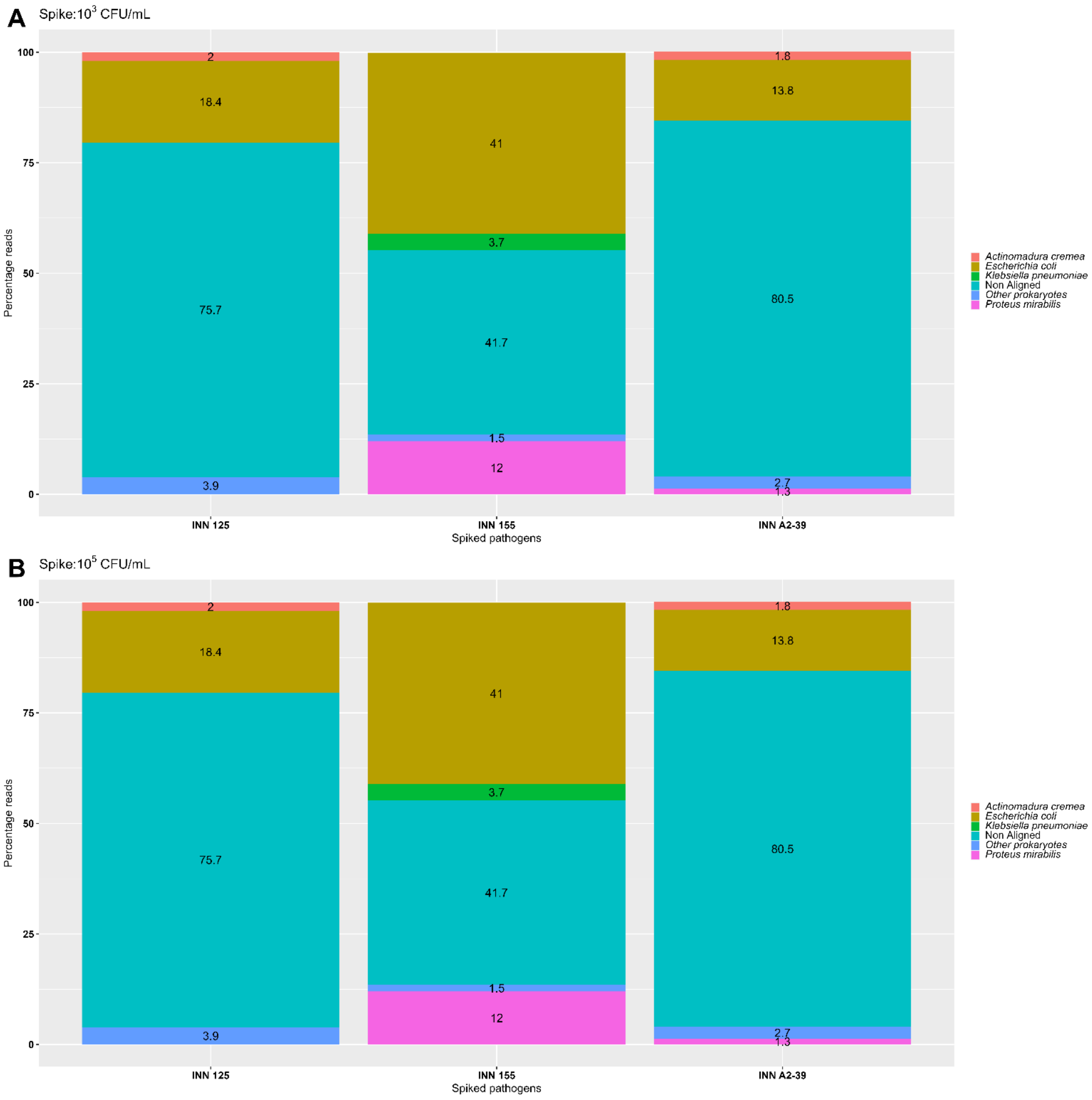


**Supplementary figure IV:** Sequencing results for the three internal test strains INN 125, INN 155 and INN A2-39. The subfigure A represents the results from the $10^3$ CFU/mL spiked samples while the subfigure B denotes the results from $10^5$ CFU/mL spiked samples. The denoted percentage reads are based on the BLAST search against the RefProk database (prokaryotic sequence data only).
