## Supplementary Figure V for "Detection of pathogens and antimicrobial resistant genes from urine within 5 hours using Nanopore sequencing"

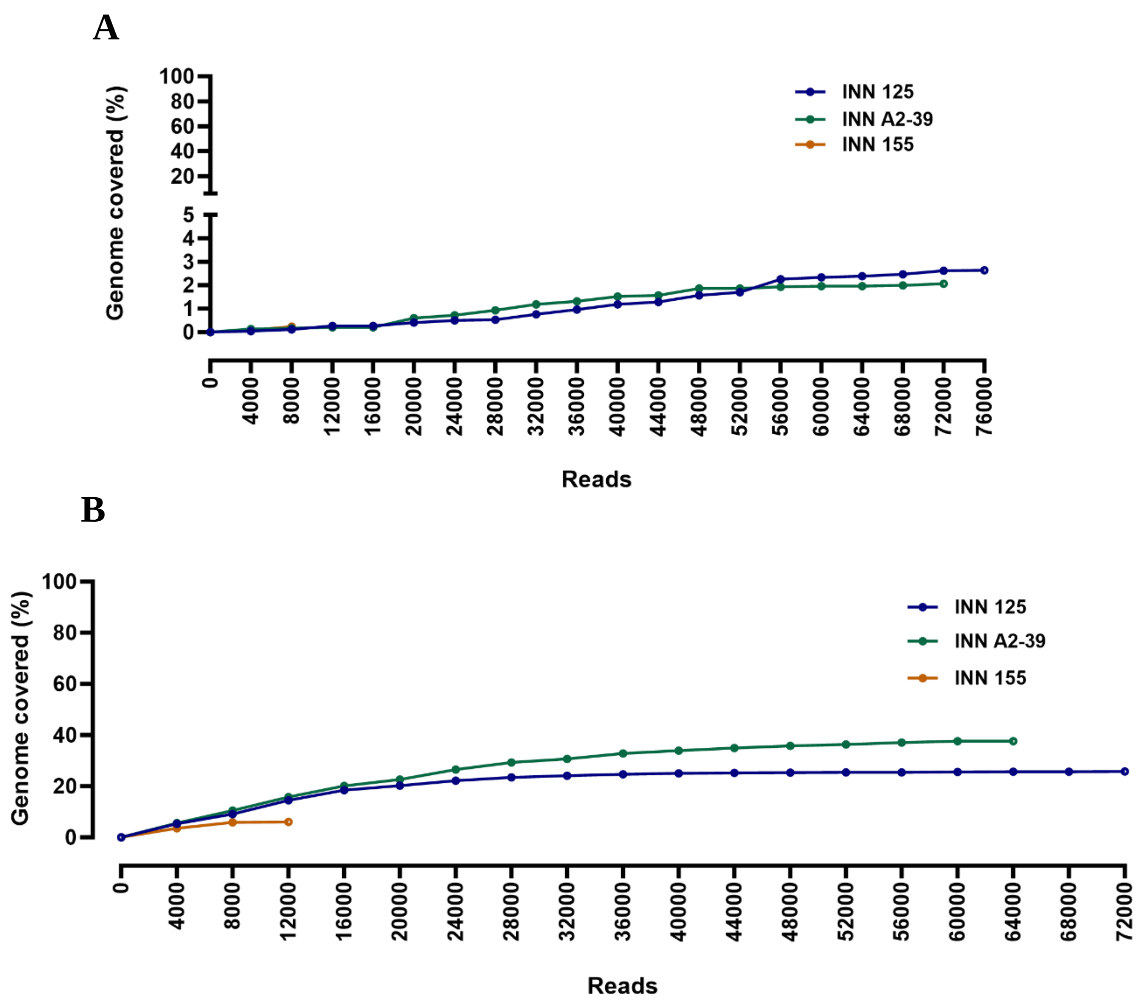


**Supplementary Figure V:** Genome coverage of the three target species INN 125, INN A2-39 and INN 155 over time. The subfigure 5. A represents the pathogen coverage obtained for the samples spiked with $10^3$ CFU/mL concentration, while the subfigure 5. B. is the pathogen coverage from the samples spiked with a pathogen concentration of $10^5$ CFU/mL.
