## Supplementary Table I for "Detection of pathogens and antimicrobial resistant genes from urine within 5 hours using Nanopore sequencing"

**Supplementary Table I:** The list of the relevant pathogenic strains and their corresponding antibiotic resistance genes used in the study.

| Bacterial strains | Reference antibiotic resistance genes (ARG's) |
|---|---|
| *E. coli* NCTC 13441 | *CTX-M-15* |
| INN 125 | *TEM-1* |
| INN A2-39 | *CTX-M-2* |
| INN 155 | *TEM-1* |
| *K. pneumoniae CCUG225T* | *SHV-11* |
| *P. mirabilis CCUG2676T* | *catA*, *tet(J)* |
| *P. aeruginosa CCUG17619* | *blaOXA-396* |
| *E. faecalis CCUG 9997* | *lsa(A)*, *tet(M)* |
| *S. aureus NCTC8325* | *fosB*, *mepA* |
