## Supplementary Table II for "Detection of pathogens and antimicrobial resistant genes from urine within 5 hours using Nanopore sequencing"

**Supplementary Table II**: The correlation between the optical density (OD) and colony formation units (CFU/mL) for pathogenic strains that were used in the study.

| Pathogens | OD | CFU/ml |
|---|---|---|
| *E. coli* NCTC 13441 | 1.39 | $8.0 \times 10^8$ |
| | 1.38 | $6.0 \times 10^8$ |
| | 1.31 | $6.1 \times 10^8$ |
| *K. pneumoniae* CCUG 225T | 1.28 | $5.5 \times 10^8$ |
| | 1.25 | $4.7 \times 10^8$ |
| | 1.26 | $6.3 \times 10^8$ |
| *S. aureus* NCTC 8325 | 1.55 | $7.8 \times 10^8$ |
| | 1.38 | $7.9 \times 10^8$ |
| *S. aureus* CCUG 35600 | 1.28 | $1.0 \times 10^9$ |
| | 1.59 | $1.7 \times 10^9$ |
| *P. mirabilis* CCUG 2676T | 2.1 | $1.3 \times 10^9$ |
| | 1.83 | $1.2 \times 10^9$ |
| | 2.2 | $3.3 \times 10^9$ |
| *E. coli 125* | 2.19 | $2.8 \times 10^9$ |
| *E. coli A2-39* | 2.12 | $3.2 \times 10^9$ |
| *E. coli* 155 | 2.19 | $2.0 \times 10^9$ |
| | 2.12 | $2.8 \times 10^9$ |
| *S. aureus* CCUG17621 | 2.25 | $6.0 \times 10^8$ |
| *E. faecalis* CCUG 9997 | 1.93 | $3.5 \times 10^9$ |
| | 2.2 | $5.2 \times 10^9$ |

| | | |
|---|---|---|
| | 2.22 | $4.6 \times 10^9$ |
| *P. aeruginosa* CCUG 17619 | 1.9 | $1.5 \times 10^9$ |
| | 1.9 | $2.4 \times 10^9$ |
