## supplementary table III for "Detection of pathogens and antimicrobial resistant genes from urine within 5 hours using Nanopore sequencing"

**Supplementary table III**: List of the primers used for confirmation of the presence of the pathogenic DNA in the DNA extracted from the spiked samples prior to nanopore sequencing.

| Target Species | Target | Product size | Primer | Primer Sequence | Reference |
|---|---|---|---|---|---|
| Human | *β-Actin* | ~ 100 bp | Human-F | CGGCCTTGGAGTGTGTATTAAGTA | (1) |
| | | | Human-R | TGCAAAGAACACGGCTAAGTGT | |
| *E. coli* | *UspA* | ~ 850 bp | uspA-F | CCGATACGCTGCCAATCAGT | (1) |
| | | | uspA-R | ACGCAGACCGTAGGCCAGAT | |
| *S. aureus* | *nuc* | ~ 65 bp | Nuc-F | GGGTTGATACGCCAGAAACG | (2) |
| | | | Nuc-R | TGATGCTTCTTTGCCAAATGG | |
| *K. pneumoniae* | *Khe* | 428 bp | Khe-Kpne-F | TGATTGCATTCGCCACTGG | (3) |
| | | | Khe-Kpne-R | GGTCAACCCAACGATCCTG | |
| *P. mirabilis* | *UrePa* | 156 bp | UrePa-pmir-F | GGTGAGATTTGTATTAATGG | (4) |
| | | | UrePa-pmir-R | ATAATCTGGAAGATGACGAG | |
| *E. faecalis* | *GroEs* | 185 bp | EfGroEs-F | GGAATTGTTCTTGCATCCGT | (5) |
| | | | EfGroEs-R | ACAATTAAGTATTCTACGCC | |
| *P. aureginosa* | *phzA2* | 325bp | PA2-F | GTTTACCGACAACCTGGAA | (6) |
| | | | PA2-R | GCAATAGCCCTGCGGATAC | |

**References:**


1. Ali J, Joshi M, Ahmadi A, Str KO, Ahmad R. Increased growth temperature and vitamin B12 supplementation reduces the lag time for rapid pathogen identification in BHI agar and blood cultures [version 2; peer review: 2 approved]. 2023;

2. Ahmadi A, Khezri A, Nørstebø H, Ahmad R. A culture-, amplification-independent, and rapid method for identification of pathogens and antibiotic resistance profile in bovine mastitis milk. Frontiers in Microbiology [Internet]. 2023 [cited 2023 May 10];13. Available from: https://www.frontiersin.org/articles/10.3389/fmicb.2022.1104701

3. Hua Wu YS. Detection and Analysis of Regional Trends of Klebsiella pneumonia Causing Liver Abscess. Clin Microbiol [Internet]. 2015 [cited 2023 Apr 20];04(04). Available from:

http://www.esciencecentral.org/journals/detection-and-analysis-of-regional-trends-of-klebsiella-pneumonia-causingliver-abscess-2327-5073-1000208.php?aid=56943

4. Zhang W, Niu Z, Yin K, Liu P, Chen L. Quick identification and quantification of Proteus mirabilis by polymerase chain reaction (PCR) assays. Ann Microbiol. 2013 Jun;63(2):683–9.

5. Teng LJ, Hsueh PR, Wang YH, Lin HM, Luh KT, Ho SW. Determination of *Enterococcus faecalis groESL* Full-Length Sequence and Application for Species Identification. J Clin Microbiol. 2001 Sep;39(9):3326–31.

6. Wang C, Ye Q, Jiang A, Zhang J, Shang Y, Li F, et al. Pseudomonas aeruginosa Detection Using Conventional PCR and Quantitative Real-Time PCR Based on Species-Specific Novel Gene Targets Identified by Pangenome Analysis. Front Microbiol. 2022 May 4;13:820431.
